# Impact of Ghost Introgression on Coalescent-based Species Tree Inference and Estimation of Divergence Time

**DOI:** 10.1101/2022.01.11.475787

**Authors:** Xiao-Xu Pang, Da-Yong Zhang

## Abstract

The species studied in any evolutionary investigation generally constitute a very small proportion of all the species currently existing or that have gone extinct. It is therefore likely that introgression, which is widespread across the tree of life, involves “ghosts,” i.e., unsampled, unknown, or extinct lineages. However, the impact of ghost introgression on estimations of species trees has been rarely studied and is thus poorly understood. In this study, we use mathematical analysis and simulations to examine the robustness of species tree methods based on a multispecies coalescent model under gene flow sourcing from an extant or ghost lineage. We found that very low levels of extant or ghost introgression can result in anomalous gene trees (AGTs) on three-taxon rooted trees if accompanied by strong incomplete lineage sorting (ILS). In contrast, even massive introgression, with more than half of the recipient genome descending from the donor lineage, may not necessarily lead to AGTs. In cases involving an ingroup lineage (defined as one that diverged no earlier than the most basal species under investigation) acting as the donor of introgression, the time of root divergence among the investigated species was either underestimated or remained unaffected, but for the cases of outgroup ghost lineages acting as donors, the divergence time was generally overestimated. Under many conditions of ingroup introgression, the stronger the ILS was, the higher was the accuracy of estimating the time of root divergence, although the topology of the species tree is more prone to be biased by the effect of introgression.

## Introduction

The development of new sequencing technologies has resulted in large datasets (for a highly limited exception, see Telford et al. 2015) that researchers have been using to reconstruct the phylogenies of species. Evolutionary trees from different genes are known to often have conflicting branching patterns, leading to ambiguous relationships between the relevant species (Maddison 1997; Nichols 2001; Degnan et al. 2009). Therefore it is important to understand the natural evolutionary processes responsible for gene tree discordance (Maddison 1997; Nichols 2001), among which incomplete lineage sorting (ILS), or the failure of two sequences from two species to coalesce in the most common ancestral species, has received the most attention (for the latest reviews, see Jiao et al. 2021; Mirarab et al. 2021).

The multispecies coalescent (MSC) model (Rannala and Yang 2003) uses a probabilistic framework to model gene tree discordance due to ILS. Coalescent-based methods are now widely used to infer the evolutionary history of species across the tree of life in case of ILS, including summary methods based on gene tree topology and full-likelihood methods based on multilocus sequence data. One class of summary methods (Larget et al. 2010; Liu et al. 2010; Mirarab et al. 2014) takes advantage of a favorable theoretical property whereby for rooted three-taxon or unrooted four-taxon trees under the MSC model, the most probable gene tree topology is the one that is concordant with the species tree (Degnan et al. 2009; Degnan 2013); if not, it is called an anomalous gene tree (AGT). ASTRAL (Mirarab et al. 2014) has arguably been the most successful of these methods owing to its high accuracy (Giarla and Esselstyn 2015; Molloy and Warnow 2018) and scalability (Mirarab et al. 2016; Yin et al. 2019). Full-likelihood methods are mainly used under the Bayesian framework, and use both gene tree topologies and branch lengths to estimate species trees as well as such parameters of population demographics as population sizes and divergence times. Programs such as *BEAST (Heled and Drummond 2010; Ogilvie et al. 2017) and BPP (Rannala and Yang 2017) are popular for inferring species tree topologies and dating speciation events (Pierron et al. 2017; Wu et al. 2018; Smith et al. 2019; Tiley et al. 2020).

Interspecific gene flow (or simply, introgression) is another major evolutionary process that can generate gene tree discordance (Slatkin and Maddison 1989). When speciation events occur close together in time, especially during rapid radiations, introgression is highly likely to occur at the same time among nonsister species as well as between sister species. Thus, it is widely expected that ILS would be at play concurrently with introgression, and this has been supported by quickly accumulating empirical evidence in recent years, such as that reviewed by Mallet et al. (2016), Taylor and Larson (2019), and Edelman and Mallet (2021). Arguably, introgression often plays no less of an important role in speciation, evolutionary innovations, and adaption than ILS. As Edelman and Mallet (2021, p.271) put it, “…, phylogenies with no evidence of gene flow are beginning to seem like the exception rather than the rule.” However, despite considerable work on species tree methods contending with ILS, only a small amount of preliminary research has incorporated cross-species gene flow into the MSC model (Elworth et al. 2019; Hibbins and Hahn 2021; Jiao et al. 2021). Worse still, current phylogenetic methods explicitly dealing with these two processes suffer from the limited number of taxa and loci that have been analyzed, largely owing to the heavy computational burden involved (Degnan 2018; Elworth et al. 2019; Blair and Ané 2020).

In this study, we consider the impacts of cross-species gene flow on estimations made according to currently available coalescent-based methods of the species tree topology and the time of divergence. In past work, Leache et al. (2014) simulated continuous cross-species migration under the model of isolation with migration (IM) (Hey 2010), and found that gene flow between nonsister species might lead to mistaken inferences regarding topology, while gene flow between sister species actually made such inference easier, both when using the summary method MP-EST (Liu et al. 2010) and full-likelihood methods. Moreover, they showed that cross-species gene flow generally results in an underestimation of the divergence times of species. By assuming episodic introgression at particular times, Solis-Lemus et al. (2016) calculated the probabilities of gene tree topologies occurring under the model of multispecies coalescent with introgression (MSci) (Yu et al. 2012), and demonstrated that popular species tree methods may reconstruct the wrong evolutionary history of species. Long and Kubatko (2018) found that AGTs might still appear under either an IM or an MSci model in case of gene flow between ancient sister species in the three-taxon cases. Jiao et al. (2020) examined the levels of gene flow between nonsister species required for incorrect estimations of the species tree, with three species subject to either a unidirectional episodic introgression or continuous migration. Although theoretical studies have clearly shown that a very low level of gene flow can lead to AGTs and, hence, mistaken species tree topologies (Jiao et al. 2020 and this study), the prevailing view among phylogeneticists is still that species tree methods using the multispecies coalescent are robust to low levels of introgression (Esquerré et al. 2021), which justifies its wide use in practice (e.g., Edwards et al. 2016). Furthermore, evolutionary studies are necessarily restricted to a subset of species or populations because the overwhelming majority of lineages that have ever lived are now extinct, and some lineages may be unsampled owing to technical limitations, or because they are not being relevant to the question at hand, or are simply unknown. It is natural that gene flow involves not only the species sampled but also “ghost” (extinct or unsampled) lineages (Eaton and Ree 2013; Pease and Hahn 2015; Hibbins and Hahn 2021; Tricou et al. 2021). However, none of the aforementioned studies has considered in sufficient details the scenarios of gene flow involving ghost lineages (for a highly limited exception, see Hibbins and Hahn 2021), nor have most empirical studies on gene flow. In addition, these studies have focused more on the impacts of gene flow on topological inference and less on its effects on estimation of divergence times.

We focus in this study on the case of three species and consider several scenarios of introgression occurring within the species tree ((*A*, *B*), *C*) under the MSci model, especially ghost introgression. We define the true species tree as one that always constitutes the true branching orders of species, regardless of whether the probability of introgression is below or above 0.5. Such a definition of a true species tree is necessary in biology because an increasing number of studies have showed that introgression can influence a majority of the genome in many groups of species across the tree of life (Fontaine et al. 2015; Thawornwattana et al. 2018; Forsythe et al. 2020; Jiao et al. 2020; Zhang et al. 2021). For each introgression scenario, we first mathematically derive explicit conditions under which AGTs exist, and then use simulations to examine the behaviors of the representative summary method, ASTRAL, and the representative full-likelihood method *BEAST. We finally quantify the impacts of gene flow among species, especially involving ghost lineages as donors of introgression, on estimations of species tree topologies and their divergence times by these methods.

## Introgression-induced formation of anomaly zones for three taxa

We focused on the case of three species with a starting species tree ((*A, B*), *C*), and then added introgression edges to it. The true species tree is always the starting tree. The data consisted of multiple loci, with three sequences at each locus (*a*, *b*, *c*)—one from each of the three species. The possible gene tree topologies considered at each locus, if introgression was lacking, were *G*_1_ = ((*a, b*), *c*), matching the species tree and occurring with the highest probability, and *G*_2_ = ((*b, c*), *a*) and *G*_3_ = ((*a, c*), *b*), not matching with the species tree and occurring with equal probabilities (Pamilo and Nei 1988). We derived the probability distributions of three gene tree topologies and identified AGT-forming regions of the parameter space. We describe several introgression scenarios below that were considered in the rooted triple, including ghost introgression, introgression between nonsister species, and introgression between extant sister species or ancestral sister species.

### Introgression from an Outgroup Ghost

Figure 1 illustrates three scenarios of introgression from an outgroup ghost that diverged earlier in time than the basal species *C*. Here, and in the sections below, we do not consider cases of ghost lineages acting as the recipient of introgression because under such conditions, a ghost lineage has no way of impacting the coalescent process of sequences (alleles) from the sampled species (Hibbins and Hahn 2021).

**Figure 1.**
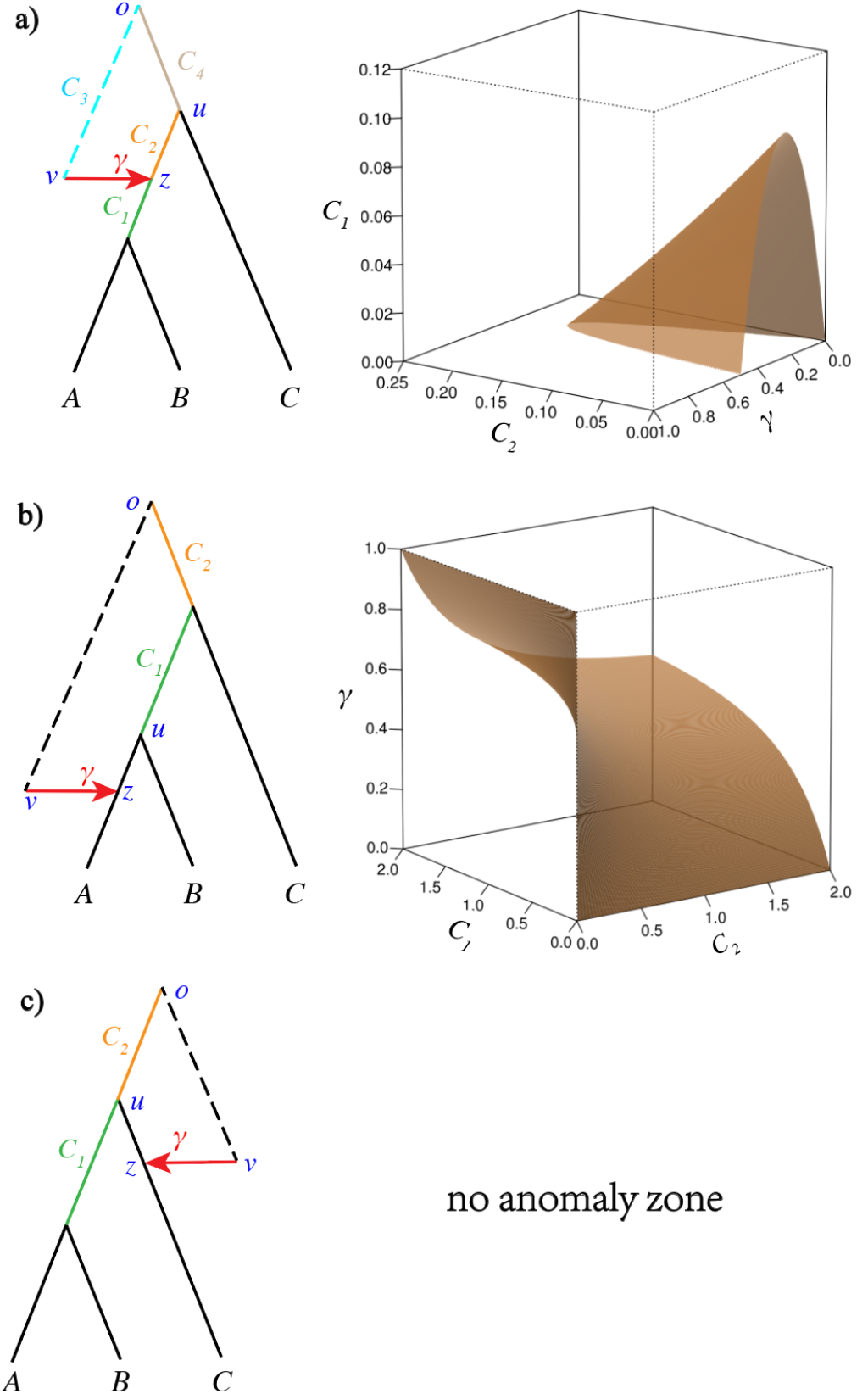
Respective anomaly zones for various patterns of introgression from an outgroup ghost. The dotted lines represent the ghost lineages. The hybrid node is labeled with the letter *z*, with the alleles in *z* having probabilities *γ* and 1-*γ* of being traced back to *v* and *u*, respectively, whereas the edge of the tree is marked with *zu* and the introgression edge with *zvo*. a) Introgression from an outgroup ghost to the ancestor of the sister species A and B. Five numerical parameters were considered, (*γ*, *C*_1_, *C*_2_, *C*_3_, *C*_4_), with branch lengths *C_i_* in coalescent units. Assuming a constant population size such that *C*_3_ = *C*_2_ + *C*_4_ and fixing *C*_4_ to 1.5, the parameter space was partitioned according to the probabilities of topologies of the gene tree. The region below the yellow surface, i.e., *P*(*G*_1_ = ((*a,b*), *c*)) < 1/3, corresponds to the anomaly zone. b) Introgression from an outgroup ghost to one of the sister species. Here, three numerical parameters were considered: (*γ*, *C*_1_, *C*_2_). The parameter space was partitioned according to the probabilities of the topologies of the gene tree. The region above the yellow surface, i.e., *P*(*G*_1_ = ((*a, b*), *c*)) < *P*(*G*_2_ = ((*b, c*), *a*)), corresponds to the anomaly zone. c) Introgression from an outgroup ghost to species C, with no possibility of anomalous gene trees.

We first analyzed the scenario with introgression from an outgroup ghost to the ancestor of A and B (Fig. 1a). Five numerical parameters, (*γ, C*_1_, *C*_2_, *C*_3_, *C*_4_), were considered, with branch lengths *C_i_* in coalescent units. Traced back in time, sequences a and b could coalesce along the branch of length *C*_1_ with a probability of 1 – *e*^−*C*_1_^. In case a and b did coalesce before the hybrid node *z* (with probability *e*^−*C*_1_^), three options were considered: (1) both a and b followed the parental path *z* → *u* with a probability of (1 – *γ*)^2^, and then had a probability 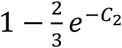 of coalescing first; (2) both followed the parental path *z* → *v* with probability *γ*^2^, then had a probability 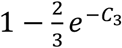 of coalescing first; and (3) one followed the path *z* → *u* but the other followed the path *z* → *v* with a probability 2*γ*(1 – *γ*), and then had a probability 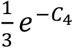 of coalescing first. In addition, *G*_2_ and *G*_3_ occurred with equal probability. The probabilities of the three gene tree topologies were derived using the equations

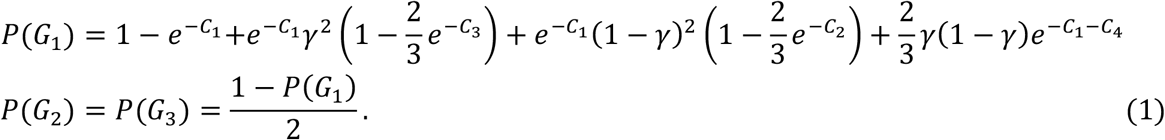

Thus, if *P*(*G*_1_) < 1/3, the occurrences of *G*_2_ and *G*_3_ were both necessarily more probable than that of *G*_1_, which matched the species tree (Fig. 1a). The fact that AGTs can exist at all under this introgression scenario may seem surprising. After all, removing either parental edge (namely, the introgression edge *vzo* and the tree edge *uz*) of the hybrid node gives rise to a tree with the topology ((*A*, *B*), *C*). Furthermore, we noted three particular characteristics of the zone of anomaly. First, 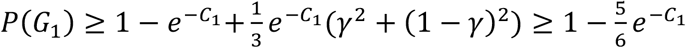, meaning that only when *C*_1_ < 0.224 could any AGT obtain. Moreover, assuming that a constant population size led to *C*_3_ = *C*_2_ + *C*_4_, then 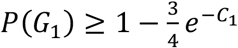 (a proof is given in the Appendix), which means that only *C*_1_ < 0.118 could produce AGTs. Second, when *C*_3_ ≥ *C*_1_, a value of *γ* greater than or equal to 0.5 would necessarily result in *P*(*G*_1_) ≥ 1/3 (see Appendix), that is, no AGTs occurred. Third, too small a probability of introgression *γ* could not result in AGTs.

For the scenario with introgression from an outgroup ghost to one sister species, i.e., A in Figure 1b, three numerical parameters, (*γ*, *C*_1_, *C*_2_), were considered. When traced back in time, if the alleles in *z* followed the parental path *z* → *u* with a probability of 1 – *γ*, and the sequences a and b coalesced along the branch of length *C*_1_ with a probability of 1 – *e*^−*C*_1_^, then the gene tree would be *G*_1_ = ((*a, b*), *c*). If the alleles in *z* followed the path *z* → *v* into the ghost lineage with a probability *γ*, and sequences b and c coalesced along the branch of length *C*_2_ with a probability 1 – *e*^−*C*_2_^, then the gene tree would be *G*_2_ = ((*b, c*), *a*). Otherwise, if both coalescence events for the three sequences occurred in the same ancestral species, then the three gene tree topologies would occur with equal probably. The probabilities of the three gene tree topologies may be described by using the equations:

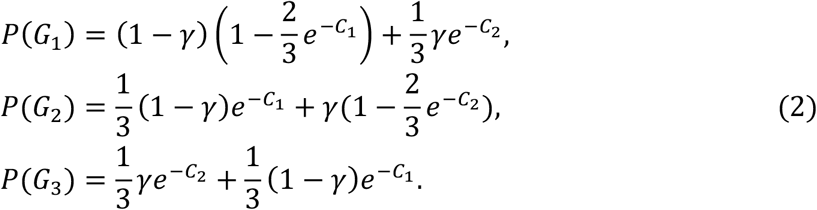

Thus, *P*(*G*_1_) < *P*(*G*_2_) corresponded to the parameter space satisfying the condition for the zone of anomaly, or *γ* > (1 – *e*^−*C*_1_^)/(2 – *e*^−*C*_1_^ – *e*^−*C*_2_^) was required for AGTs. Clearly, the smaller the value of *C*_1_ was and, hence, the stronger ILS was, the smaller was the probability of introgression *γ* required for AGTs to occur. In other words, a more ILS necessitates less introgression to give rise to AGTs. In the extreme case of a very strong ILS with *C*_1_ approaching zero, an arbitrarily low value of *γ* would be sufficient for AGTs to obtain. By contrast, we note that even *γ >* 0.5 might not necessarily have led to AGTs under some conditions, such as when *C*_1_ > *C*_2_.

In case of introgression from an outgroup ghost to the basal species C (Fig. 1c), three numerical parameters, (*γ, C*_1_, *C*_2_), were considered. When traced back in time, if the sequence c followed the path *z* → *u* with a probability of 1 – *γ*, the sequences a and b could coalesce along the branch of length *C*_1_ with probability 1 – *e*^−*C*_1_^. If sequence c followed the path *z* → *v* with probability *γ*, the sequences a and b could coalesce along the branch of length *C*_1_ + *C*_2_ with probability 1 – *e*^−(*C*_1_+*C*_2_^). Thus, in this condition, *G*_2_ and *G*_3_ occurred with equal probability, and the probability of *G*_1_ is given by the equation:

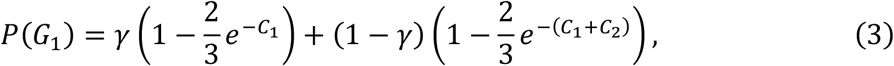

which is always larger than 1/3. In other words, anomalous gene trees are impossible in this condition.

### Introgression between Nonsister Species, and some related scenarios of Ingroup Ghost Introgression

In this subsection, we follow Jiao et al. (2020) by using “inflow” and “outflow” to denote different directions of introgression: “Inflow” refers to the direction from the basal species C to either sister species (namely, B in Fig. 2a), and “outflow” denotes the opposite direction of introgression (Fig. 2b). We also considered two similar scenarios of introgression; One was from a C-derived ingroup ghost (i.e., with the ghost lineage speciating after the appearance of the basal species C) to one sister species (as illustrated in Fig. 2c), and other was from a sister species-derived ingroup ghost to C (Fig. 2d). For all four cases, three numerical parameters were considered: (*γ*, *C*_1_ *C*_2_). When traced back in time, if the alleles in *z* followed the path *z* → *u* with a probability of 1 – *γ*, and sequences a and b coalesced along the branch of length *C*_2_ with a probability of 1 – *e*^−*C*_2_^, the gene tree were *G*_1_ = ((*a, b*), *c*). If the alleles in *z* followed the path *z* → *v* with a probability of *γ*, and sequences b and c coalesced along the branch of length *C*_1_ with a probability of 1 – *e*^−*C*_1_^, the gene tree was *G*_2_ = (*b, c*), *a*). Otherwise, if both coalescence events for the three sequences occurred in the same ancestral species, the three gene tree topologies occurred with equal probability. These four scenarios yielded the same mathematical expressions for the probabilities of the three gene tree topologies, as given by Jiao et al. (2020), without considering ghost introgressions:

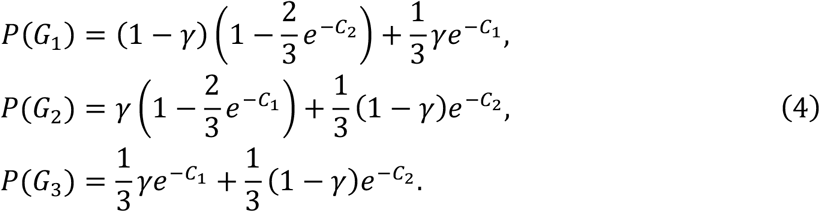

**Figure 2.**
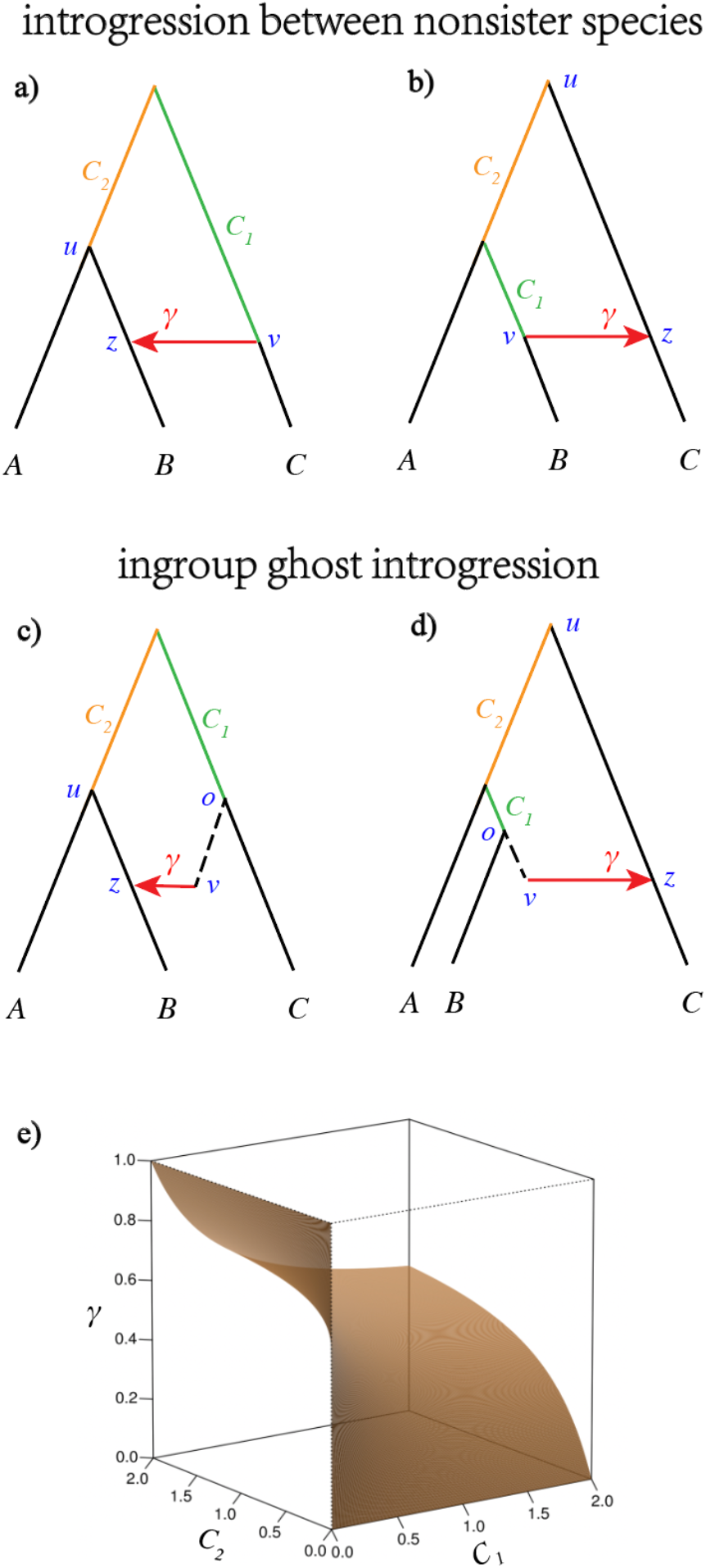
Respective anomaly zones for introgression between nonsister species as well as some scenarios of ingroup ghost introgression. (a, b) Scenarios of introgression between nonsister species with (a) the direction of inflow *C* → *B* and (b) direction of outflow *B* → *C*. (c, d) Scenarios of introgression (c) from a *C*-derived ingroup ghost to *B* and (d) from a *B*-derived ingroup ghost to *C*. For all four introgression scenarios, three numerical parameters were considered, namely, (*γ*, *C*_1_, *C*_2_), with branch lengths *C*_1_ and *C*_2_ in coalescent units. Note that *C*_1_ represents different branches in the four scenarios. The hybrid node in each case is labeled with the letter *z*, with the alleles in *z* having the probabilities *γ* and 1 – *γ* of being traced back to *v* and *u*, respectively, whereas the edge of the tree is marked with *zu* and the introgression edge with *zv* or *zvo*. (e) The parameter space was partitioned according to the probabilities of gene tree topologies in the above four scenarios. The region above the yellow surface, i.e., *P*(*G*_1_ = ((*a, b*), *c*)) < *P*(*G*_2_ = ((*b, c*), *a*)), corresponds to the anomaly zone.

The anomalous zone corresponded to the parameter space satisfying the condition *γ* > (1 – *e*^−*C*_2_^)/(2 – *e*^−*C*_2_^ – *e*^−*C*_2_^) (Fig. 2e). Other things being equal, the smaller the value of *C*_2_ was, the smaller was the amount of introgression required for AGT to occur, indicating a strong interaction between introgression and ILS in the possible appearance of AGTs. When ILS was very strong with an exceedingly small *C*_2_, even an arbitrarily small *γ* was sufficient to result in AGT. However, we found that a high probability of introgression of *γ* > 0.5 did not necessarily lead to AGT, as already shown for outgroup ghost introgression (Fig. 1b). The exception to all this was as follows: For inflow introgression, *C*_1_ > *C*_2_ when assuming a constant population size such that a high probability of introgression of *γ* > 0.5 inevitably led to AGT. Under equivalent conditions, outflow introgression required a higher level of introgression for AGT to occur.

### Introgression between Ancestral Sister Species, and some related scenarios of Ingroup Ghost Introgression

Figure 3 illustrates scenarios of introgression between ancestral sister lineages with different directions of introgression, together with similar cases of ingroup ghost introgression. We first considered the scenario of introgression from a C-derived ingroup ghost to the ancestor of sister species (Fig. 3c), with five numerical parameters considered: (*γ, C*_1_, *C*_2_, *C*_3_, *C*_4_). When traced back in time, sequences a and b would have coalesced along the branch of length *C*_1_ with a probability of 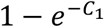. In case a and b did not coalesce before the hybrid node *z*, three options were considered: (1) Both of them followed the path *z* → *u* with a probability of (1 – *γ*)^2^, and then had a probability of 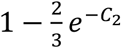 of coalescing; (2) both of them followed the path *z* → *v* with a probability of *γ*^2^, and then had a probability of 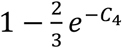 of coalescing; or (3) one of them followed the path *z* → *u* but the other followed the path *z* → *v* with a probability of 2*γ*(1 – *γ*), and then they had a probability of 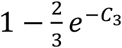 of coalescing. Thus, in this condition, *G*_2_ and *G*_3_ occurred with equal probability, and the probabilities of occurrence of the three gene tree topologies were given by the equations

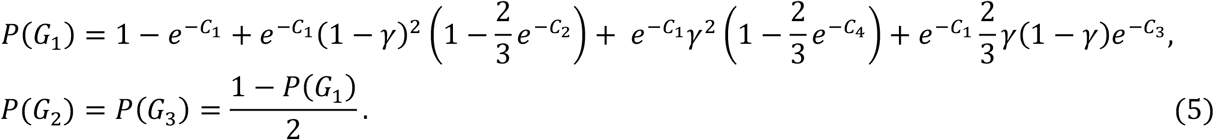

**Figure 3.**
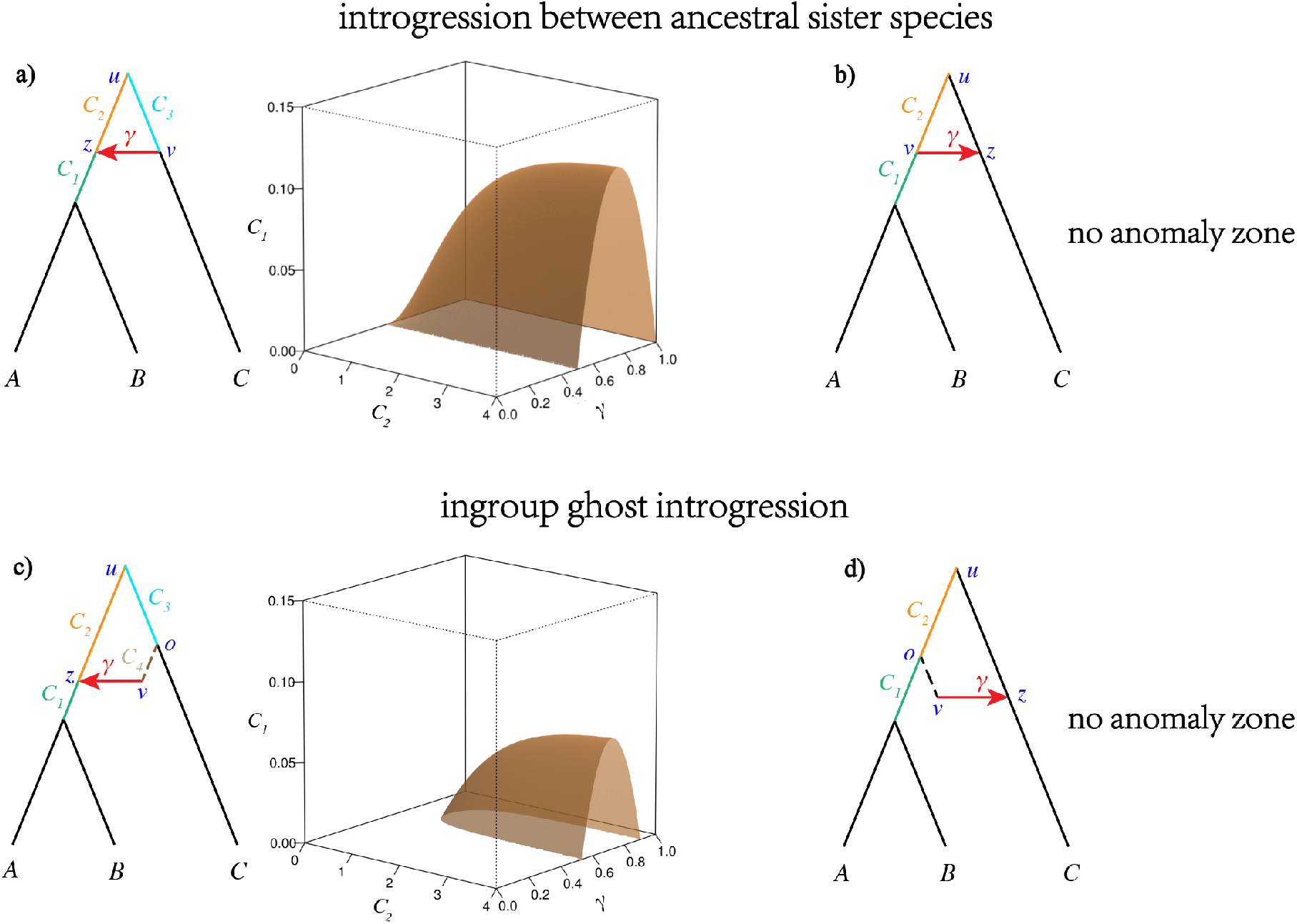
Respective anomaly zones for different scenarios of introgression between ancestral sister species as well as some cases of ingroup ghost introgression. The hybrid node in each case is labeled with the letter *z*, with the alleles in *z* having the probabilities *γ* and 1 – *γ* of being traced back to *v* and *u*, respectively, whereas the edge of the tree is labeled with *zu* and the introgression edge with *zv* or *zvo*. a) Introgression between ancestral sister species with species C as the donor of gene flow. Here, four numerical parameters were considered, namely, (*γ*, *C*_1_, *C*_2_, *C*_3_), with branch lengths *C_i_* in coalescent units. Assuming a constant population size such that *C*_2_ = *C*_3_, the parameter space was partitioned according to the probabilities of the gene tree topologies. The region below the yellow surface, i.e., *P*(*G*_1_ = ((*a, b*), *c*)) < 1/3, corresponds to the anomaly zone. b) Introgression between ancestral sister species with species C as the recipient of gene flow. There is no possibility for anomalous gene trees in this scenario. c) Introgression from a C-derived ingroup ghost to the ancestor of a sister species. Five numerical parameters were considered, namely, (*γ*, *C*_1_, *C*_2_, *C*_3_, *C*_4_). Assuming a constant population size such that *C*_2_ = *C*_3_ + *C*_4_ and fixing *C*_4_ to 0.1, the parameter space was partitioned according to the probabilities of the gene tree topologies. The region below the yellow surface, i.e., *P*(*G*_1_ = ((*a, b*), *c*)) <1/3, corresponds to the anomaly zone. d) Introgression from an ingroup ghost, derived from the ancestor of a sister species, to species C. There is no possibility of anomalous gene trees in this scenario.

In particular for introgression between ancestral sister species along the direction from *C* to their ancestor (Fig. 3a), the analysis by Long and Kubatko (2018) is merely a special case with *C*_4_ = 0. For both scenarios, when *P*(*G*_1_) < 1/3, *G*_2_ and *G*_3_ were AGTs, the corresponding anomaly zones were as shown on the right side of Figures 3a and 3c, respectively. Three particular features of these anomaly zones should be noted: First, 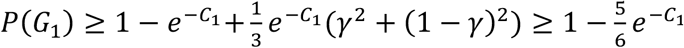, which means that only *C*_1_ < 0.224 could generate AGTs. Furthermore, if we assume a constant population size such that *C*_2_ = *C*_3_ in Figure 3a and *C*_2_ = *C*_3_ + *C*_4_ in Figure 3c, then 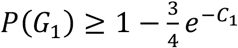, which means that only *C*_1_ < 0.118 can produce AGTs (see Appendix); second, when *C*_2_ ≥ *C*_3_, a value of *γ* less than or equal to 0.5 would have necessarily led to *P*(*G*_1_) ≥ 1/3 (see Appendix); in other words, only when a sufficiently high probability of introgression was obtained (i.e., *γ* > 0.5) could AGTs have been generated; and third, an introgression probability *γ* that was too large or too small could not produce AGTs.

In the scenario involving introgression from the ancestor of the sister species to species C (Fig. 3b), or from the ancestor-derived ghost to species C (Figs. 3d), if sequence c had followed the path *z* → *u* with a probability 1 – *γ*, sequences a and b would have been able to coalesce along the branch of length *C*_1_ + *C*_2_ with a probability of 1 – *e*^−(*C*_1_+*C*_2_)^; if sequence c had followed the path *z* → *v* with a probability of *γ*, sequences a and b would have been able to coalesce along the branch of length *C*_1_ with a probability of 1 – *e*^−*C*_1_^. Thus, for these two cases, *G*_2_ and *G*_3_ were once again equally probable, and the probability of *G*_1_ was given as:

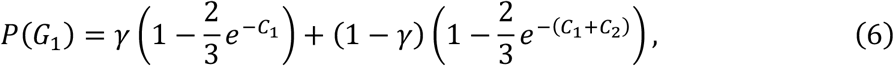

which is always larger than 1/3. Therefore, no anomaly zone was possible for these two introgression scenarios.

### Introgression between Extant Sister Species, and some related scenarios of Ingroup Ghost Introgression

Consider the scenarios of introgression between sister species and introgression from an ingroup ghost derived from A to B (Figs. 4a–b). Tracing back in time, if sequence c had followed the path *z* → *u* with a probability of 1 – *γ*, then sequences a and b would have been able to coalesce along the branch of length *C*_2_ with a probability of 1 – *e*^−*C*_2_^. If it had followed the path *z* → *v* with a probability of *γ*, then sequences a and b would have been able to coalesce along the branch of length *C*_1_ + *C*_2_ with a probability of 1 – *e*^−(*C*_1_+*C*_2_)^. Thus, in these two scenarios, *G*_2_ and *G*_3_ occurred with equal probability, and the probability of *G*_1_ can be given by the equation:

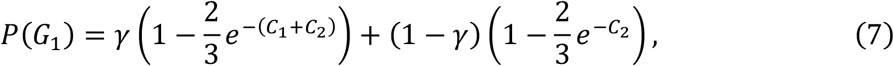

which is always larger than 1/3.

**Figure 4.**
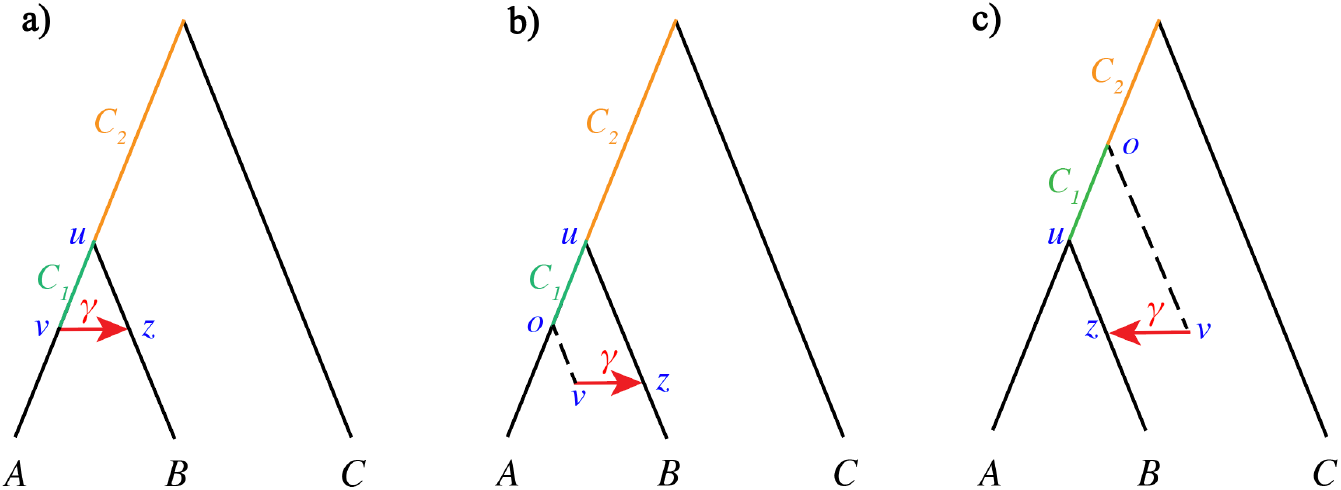
No anomaly zones are possible for the case of introgression between terminal sister species and two related cases of ingroup ghost introgression. The hybrid node in each case is labeled with the letter *z*, with the alleles in *z* having the probabilities *γ* and 1-*γ* of being traced back to *v* and *u*, respectively. Moreover, the edge of the tree is marked with *zu* and the edge of introgression with *zv* or *zvo*. a) Introgression between sister species. b) Introgression from an A-derived ingroup ghost to B. c) Introgression from an ingroup ghost, derived from the ancestor of the sister species, to B.

In the scenarios of introgression from an ingroup ghost stemming from the ancestor to one of its descendant species (Fig. 4c), if the sequence c had followed the path *z* → *u* with a probability 1 – *γ*, then sequences a and b would have been able to coalesce along the branch of length *C*_1_ + *C*_2_ with a probability of 1 – *e*^−(*C*_1_+*C*_2_)^, whereas if it had followed the path *z* → *v* with a probability of *γ*, the sequences a and b would have been able to coalesce along the branch of length *C*_2_ with a probability of 1 – *e*^−*C*_2_^. Thus, under this condition, *G*_2_ and *G*_3_ would have occurred with equal probability, and the probability of *G*_1_ would have been:

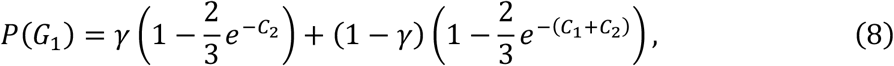

which is also always larger than 1/3. Thus, anomalous behaviors were concluded to not have existed for any of the three cases in Figure 4.

## Species tree estimation in the presence of introgression: simulations

### Data Simulations

We simulated 10 introgression scenarios to evaluate the performances of coalescent-based species tree methods. For all scenarios, the edge linking the divergence of outgroup species O to the most recent common ancestor (MRCA) of the three focal species and the ghost lineages was set to five, in coalescent units. We used the ms simulator (Hudson 1983) to simulate 1,000 gene trees, with one haploid individual per species. The simulated gene trees were used as input to simulate DNA sequences using seq-gen (Rambaut and Grass 1997), with 1,000 base pairs under a rate of population mutation of 0.036, the HKY model with a transition–transversion ratio of one, and frequencies of A, C, G, and T of 0.300414, 0.191363, 0.196748, and 0.311475, respectively. The 1,000 true gene trees and sequences of 50 loci were used as inputs to ASTRAL (ASTRAL-III version) and *BEAST (StarBEAST2 version (Ogilvie et al. 2017)), respectively. We ran ASTRAL with default settings, and used species O to root the estimated unrooted species trees. We ran the *BEAST MCMC algorithm for 50 million generations, sampling every 5,000 steps together with a 10% burn-in. The HKY model of nucleotide substitution with kappa=1 was used for all loci to match the simulation conditions. A strict molecular clock was assumed, and the clock rate was fixed at one. A Yule process prior was used to estimate the divergence times in the species tree, and the model of population size was set to “constant.” The sequences A, B, and C were set to be monophyletic. Lastly, the posterior sample of the phylogenetic species trees produced by *BEAST was summarized into a maximum clade credibility tree using TreeAnnotator v2.6.3 by using the mean heights rule.

We determined the numbers of the three estimated species tree topologies ((*A, B*), *C*), (*A*, (*B, C*)), and ((*A, C*), *B*) across 20 replicates (omitting the outgroup species O for simplicity), and calculated the mean local posterior probability (local PP) (Sayyari and Mirarab 2016) provided by ASTRAL and the posterior clade probability (PP) provided by *BEAST for the clade of the inferred sister species. The two types of support values for clades also represented the confidence level for the entire inferred topology of the species tree. Furthermore, we measured errors in the estimates of divergence times *τ* for the MRCA of the three focal species in the tree, with *τ* = *θt*/2 (rate of population mutation *θ* = 0.036, and time *t* in coalescent units for the MRCA of species A, B, and C).

### Results of scenarios of Outgroup Ghost Introgression

Figure 5 presents the results of the simulated scenarios of introgression from an outgroup ghost to the ancestor of the sister species. ASTRAL performed poorly in the anomaly zone, as expected. Specifically, for a ghost introgression event very rapidly followed by a lineage-splitting event that results in two sister species A and B (e.g., *t*_1_ = 0.05), ASTRAL inferred the false species trees (*A*, (*B, C*)) or ((*A, C*), *B*) at *γ* < 0.5, notably for a distant outgroup ghost (*t*_2_ > 0.5). Further, as *γ* was increased from 0 to 0.5, the accuracy of the topological inference decreased first and then increased (Fig. 5b). The local PPs of ASTRAL had a similar variational trend to that of the accuracy of the topological inference, and were found to be as low as about 0.7 when accuracy was low (Fig. 5d). *BEAST also worked poorly with small values of *t*_1_, but appeared to be insensitive to the distance of the outgroup ghost as measured by *t*_2_ (Fig. 5c). *BEAST also provided low support of about 0.7 in case of a lack of accuracy (Fig. 5d). Interestingly, the divergence times for the MRCA of the three species were overestimated, more so with a larger value of *γ* (Fig. 5e). Posterior estimates of divergence time 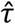 remained relatively constant, and matched the true values when *γ* was below 0.5, but then began to increase substantially despite correct topological inferences (Fig. 5e). In other words, even though the tree topology had been correctly inferred, lengths of the tree branches might still have been significantly overestimated in the presence of massive introgression.

**Figure 5.**
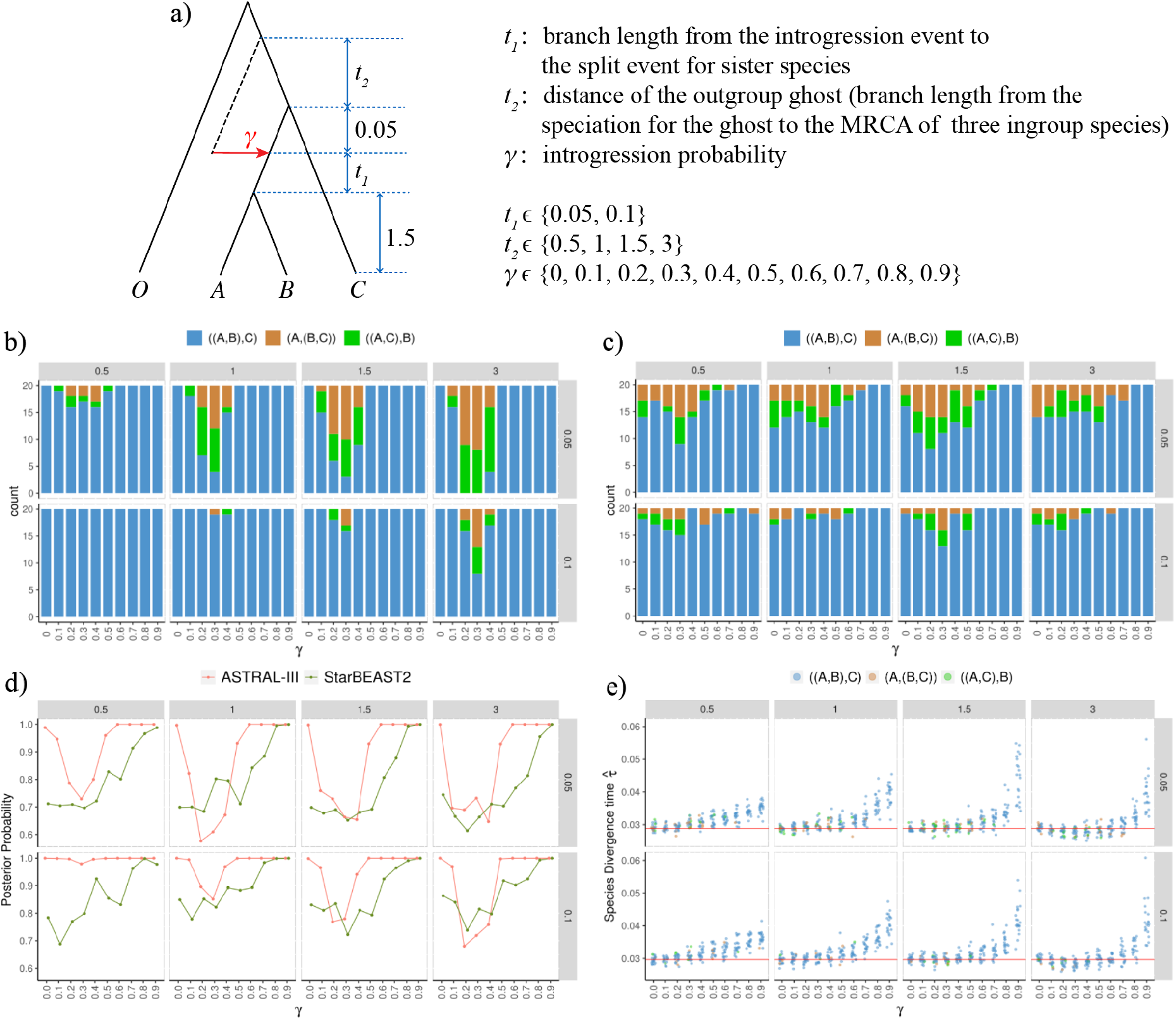
Results of simulation of introgression from an outgroup ghost to the ancestor of a sister species. a) The simulated introgression scenarios and parameter settings. The outgroup species is labeled with the letter O. The lengths of the branches *t_i_* are each in coalescent units. (b–e) Plots of the results of simulation. The numbers specified on the strips on the top and right of each plot represent *t*_2_ and *t*_1_, respectively, and *γ* is labeled on the x-axis. b) The numbers of the tree topology of three species inferred by ASTRAL-III (with the outgroup species O omitted for simplicity) among 20 replicates. Three colors, blue, orange, and green, represent the three different topologies ((A,B),C), (A,(B,C)), and ((A,C),B), respectively. c) The numbers of tree topologies of three species inferred by StarBEAST2. d) Local posterior probability (ASTRAL-III) and posterior clade probability (StarBEAST2) of the node of the sister species. Each data point represents the mean value across 20 replicates, and is jittered horizontally to avoid clutter. e) Estimated divergence time for the most recent common ancestor (MRCA) of the three ingroup species. The points and the red line represent the estimates and the true value, respectively.

In scenarios of introgression from an outgroup ghost to a sister species (Fig. 6), the accuracy of the species tree constructed by ASTRAL dropped as *t*_1_ decreased, or as *t*_2_ and *γ* increased (Fig. 6b). For a hard species tree with very short internal branch lengths at *t*_1_ = 0.05, modest introgression (e.g., *γ* = 0.1~0.2) was found to apparently cause sharp reductions in accuracy, especially for a relatively distant outgroup ghost (e.g., *t*_2_ = 0.9) (Fig. 6b). *BEAST was found to be more robust against introgression than ASTRAL in this scenario (Fig. 6c), although their qualitative behaviors were basically similar. At a high ILS with a low *t*_1_, i.e., *t*_1_ ≤ 0.1, accuracy declined markedly with increasing introgression. However, for moderate levels of ILS at *t*_1_ ≥ 0.5, the true topology ((*A, B*), *C*) was estimated with a higher accuracy and high confidence at *γ* ≤ 0.7. Note that, importantly, both ASTRAL and *BEAST were observed to provide strong support for the false topology (*A*, (*B, C*)) in certain conditions, such as for excessive introgression with *γ* > 0.7. The divergence times for the MRCA of the three focal species were overestimated, and the extent of overestimation was associated with the level of ILS (*t*_1_), the distance of the outgroup ghost (*t*_2_), and the estimated topology (Fig. 6e). Specifically, for a high ILS with *t*_1_ ≤ 0.1, *BEAST all too easily inferred the false topology (*A*, (*B, C*)), and the posterior estimates for divergence times 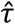 were highly overestimated at *γ* > 0.5 for a distant outgroup ghost (e.g., *t*_2_ = 0.9). By contrast, for medium levels of ILS with *t*_1_ = 0.9, the values of 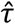 were close to those estimated with *γ* = 0 (ILS only), except for the incorrectly inferred topology at *γ* = 0.9.

**Figure 6.**
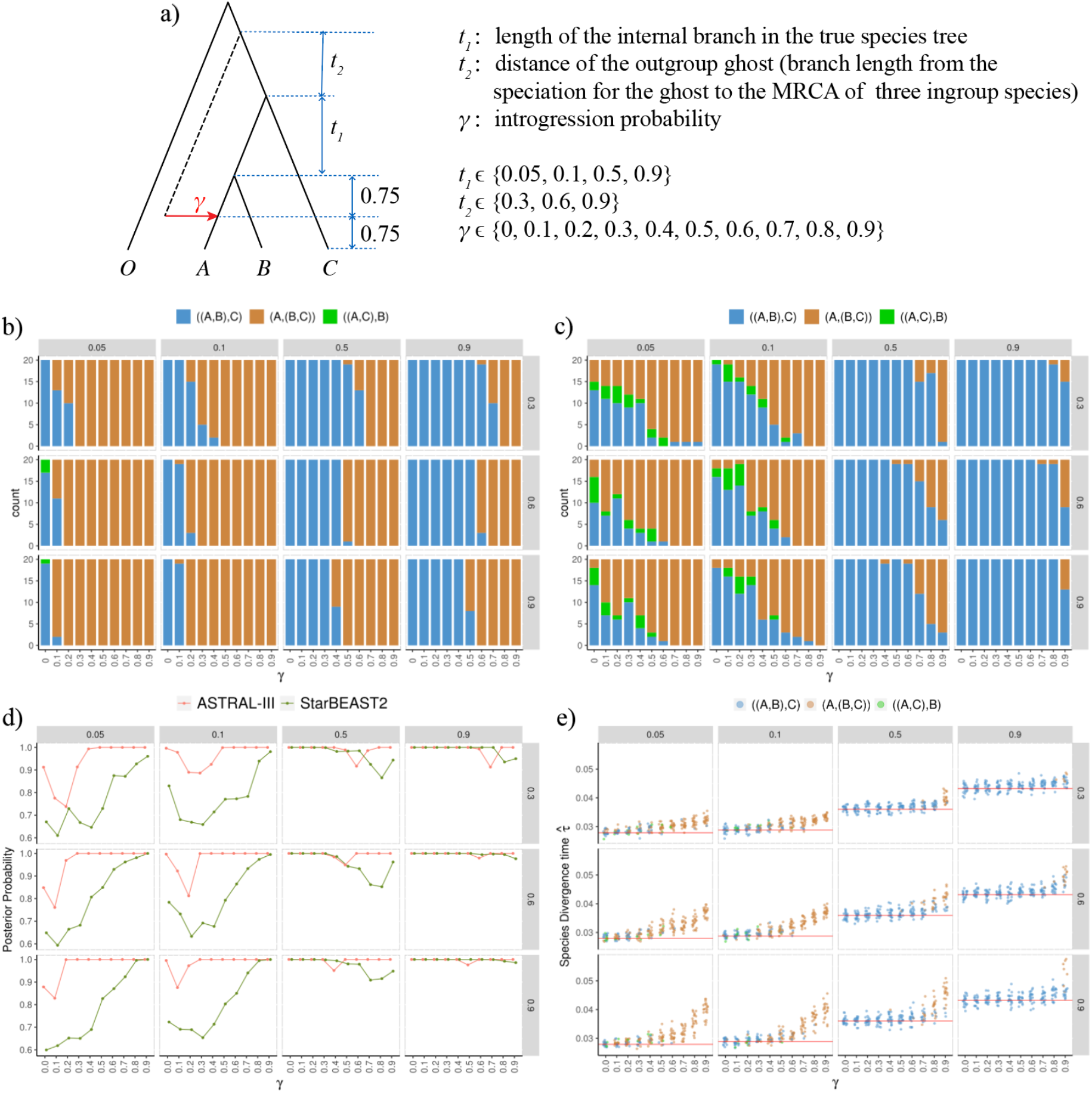
Results of simulation of introgression from an outgroup ghost to one of the sister species. a) The simulated introgression scenarios and parameter settings. The outgroup species is labeled with O. The lengths of the branches *t_i_* are in coalescent units. (b–e) Plots of the results of simulations. The numbers specified on the strips on the top and right of each plot represent *t*_1_ and *t*_2_, respectively, and *γ* is labeled on the x-axis. b) The numbers of topologies of three species trees inferred by ASTRAL-III (with the outgroup species O omitted for simplicity) among 20 replicates. Three colors, blue, orange, and green, represent the three topologies ((A,B),C), (A,(B,C)), and ((A,C),B), respectively. c) The numbers of topologies of three species trees inferred by StarBEAST2. d) Local posterior probability (ASTRAL-III) and posterior clade probability (StarBEAST2) of the node of the sister species. Each data point represents the mean value across 20 replicates, and is jittered horizontally to avoid clutter. e) Estimated divergence times for the MRCA of the three ingroup species. The points and the red line represent the estimates and the true value, respectively.

For scenarios with introgression from an outgroup ghost to species C (Fig. S1), we set a short internal branch in the species tree with *t*_1_ = 0.05 and changed the distance of the outgroup ghost *t*_2_. ASTRAL inferred the species tree topology highly accurately and local PPs increased with *γ*. The accuracy and PPs of *BEAST also increased with *γ*. Overall, the results showed that introgression in this condition had a similar synergistic effect to that between sister species (Leache et al. 2014). The divergence times of the MRCA of the three focal species had been overestimated (Fig. S1e). When the distance of the outgroup ghost *t*_2_ was not short (e.g., *t*_2_ = 1), the posterior estimates for divergence times 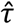 were accurate at *γ* < 0.5 but were severely overestimated at *γ* ≥ 0.5.

### Results of scenarios of Introgression between Nonsister Species, and some Related Ingroup Ghost Introgressions

Figures 7 and 8 summarize the results of using the ASTRAL and *BEAST methods for scenarios of introgression between nonsister species as well as some related ingroup ghost introgression scenarios. Note that the inflow introgression *C* → B (the left side of Fig. 7a) and introgression from a C-derived ingroup ghost to B (the right side of Fig. 7a) generated the same distribution of gene trees when the parameters were as annotated as in the figures. In other words, the two scenarios yielded the same results in the estimated species tree topology and divergence time. Similarly, for the outflow introgression (*B* → C) (the left side of Fig. 8a) and the introgression from a B-derived ingroup ghost to C (the right side of Fig. 8a), the conclusions of one scenario can be applied to the other. In particular, in addition to the parameter settings for introgression from a C-derived ingroup ghost to B in Figure 7, we explored the behaviors of coalescent-based methods when the divergence time of the ghost predated the time of splitting of the sister species in Figure S2.

**Figure 7.**
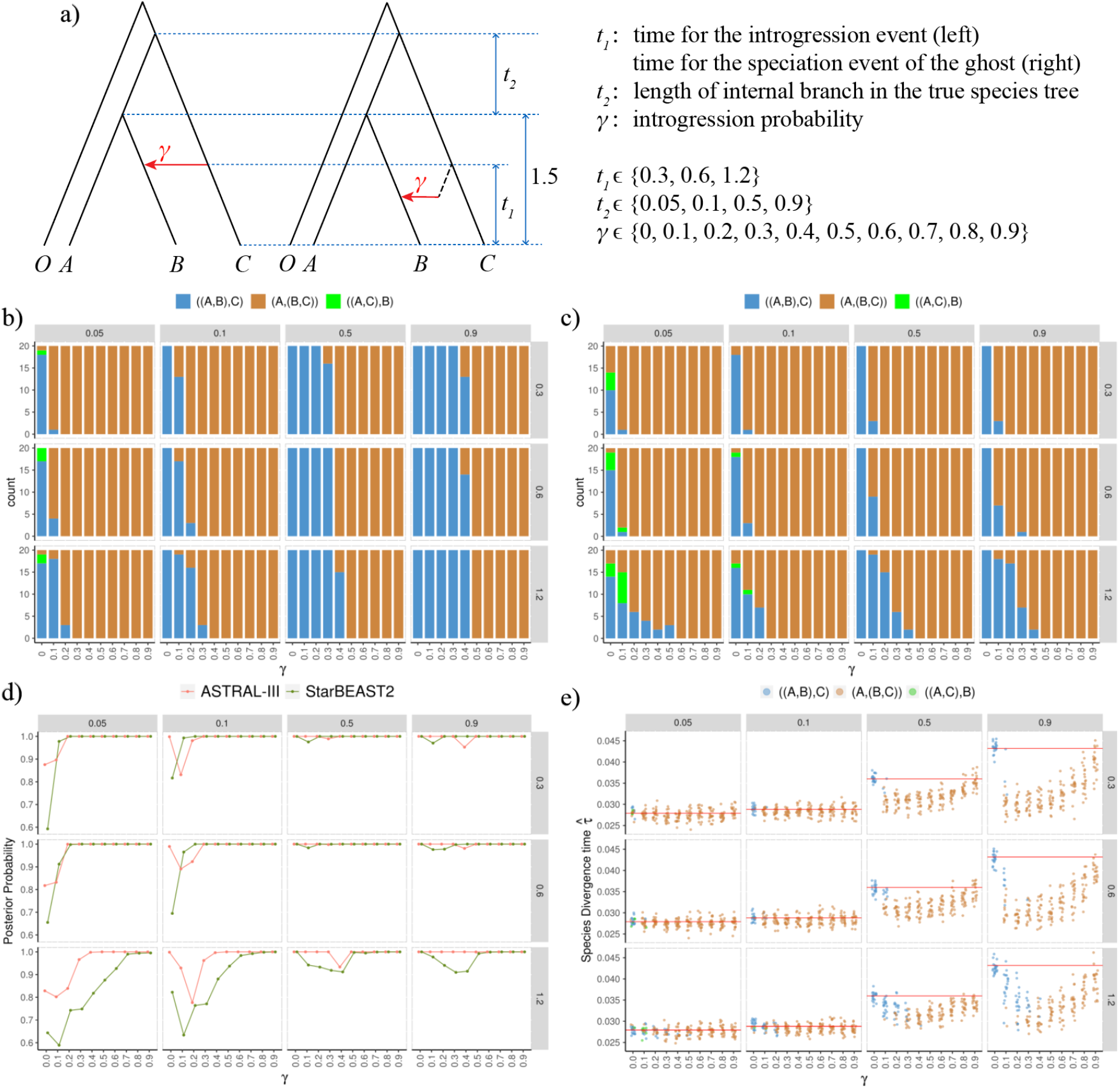
Results of simulation of introgression between nonsister species with inflow direction *C* → *B*, as well as introgression from a C-derived ingroup ghost to B. a) The simulated introgression scenarios and parameter settings. The outgroup species in each case is labeled with O. The lengths of the branches *t_i_* are in coalescent units. The two scenarios generated the same distribution of gene trees with coalescent times under the same parameter settings (*t*_1_, *t*_2_, *γ*) when one individual was sampled per species. Thus, they had the same results in our simulation. (b–e) Plots of the results of simulations. The numbers specified on the strips on the top and right of each plot represent *t*_2_ and *t*_1_, respectively, and *γ* is labeled on the x-axis. b) The numbers of topologies of three species trees inferred by ASTRAL-III (with the outgroup species O omitted for simplicity) among 20 replicates. c) The numbers of topologies of three species trees inferred by StarBEAST2. d) Local posterior probability (ASTRAL-III) and posterior clade probability (StarBEAST2) of the node of the sister species. Each data point represents the mean value across 20 replicates, and is jittered horizontally to avoid clutter. e) Estimated divergence times for the MRCA of the three species. The points and the red line represent the estimates and the true value, respectively.

**Figure 8.**
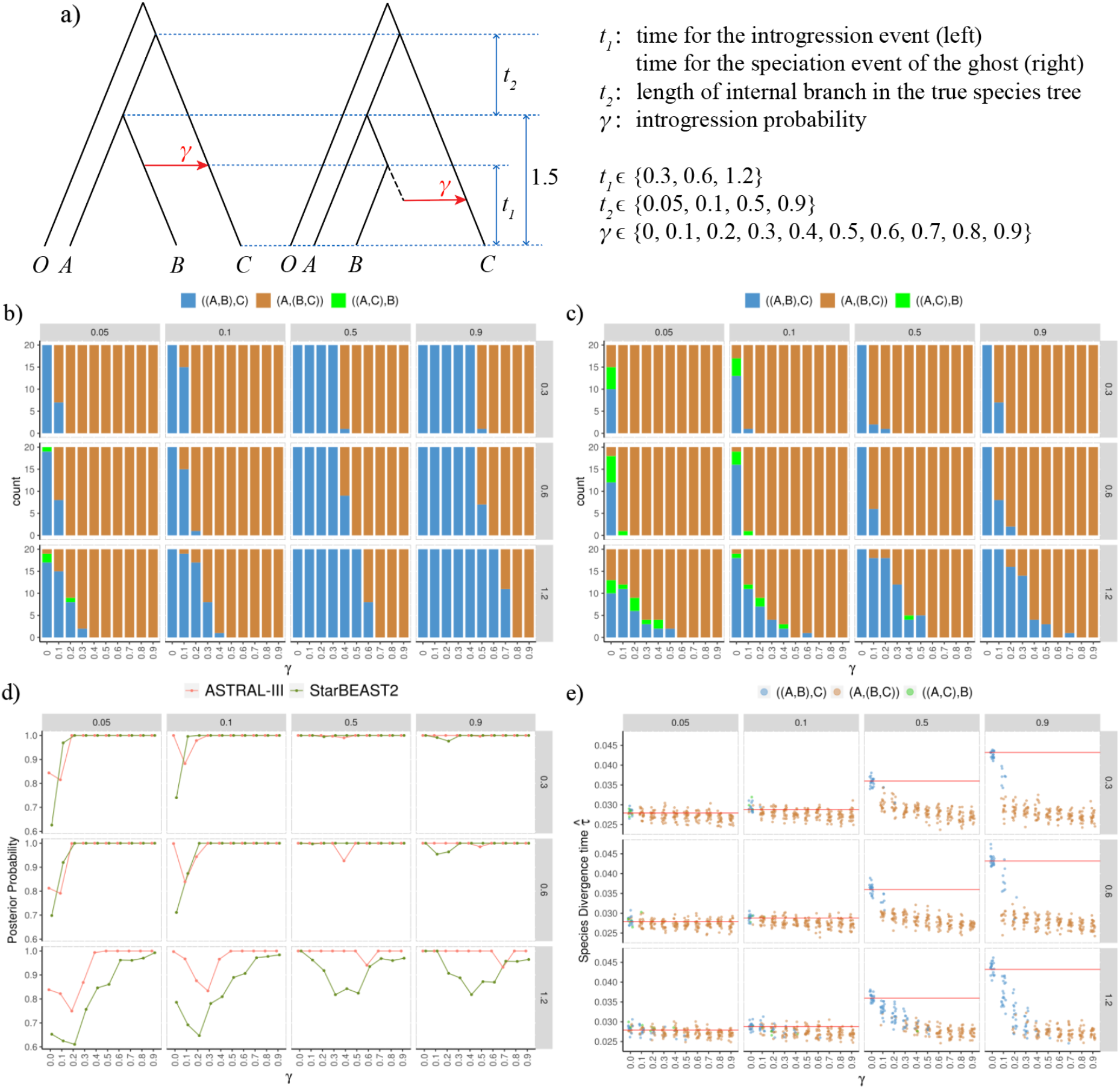
Results of simulation of introgression between nonsister species with outflow direction *B* → *C*, as well as introgression from a B-derived ingroup ghost to C. a) The simulated introgression scenarios and parameter settings. The outgroup species in each case is labeled with O. The lengths of the branches *t_i_* are in coalescent units. The two scenarios generated the same distribution of gene trees with coalescent times under the same parameter settings (*t*_1_, *t*_2_, *γ*) when one individual was sampled per species, so they share the same results in our simulation. (b–e) Plots of the results of simulations. The numbers specified on the strips on the top and right of each plot represent *t*_2_ and *t*_1_, respectively, and *γ* is labeled on the x-axis. b) The numbers of topologies of three species trees inferred by ASTRAL-III (with the outgroup species O omitted for simplicity) among 20 replicates. c) The numbers of topologies of three species trees inferred by StarBEAST2. d) Local posterior probability (ASTRAL-III) and posterior clade probability (StarBEAST2) of the node of the sister species. Each data point represents the mean value across 20 replicates, and is jittered horizontally to avoid clutter. e) Estimated divergence times of the MRCA of the three ingroup species. The points and the red line represent the estimates and the true value, respectively.

For both methods, the accuracy of the estimated species tree topology was found to decrease as the levels of ILS (*t*_2_) and introgression (*γ*) increased, and as *t*_1_ approached the splitting node of species A and B (Figs. 7b and c, 8b and c). In particular for the prevalent ILS with *t*_2_ ≤ 0.1, the introduction of even a modest introgression, such as *γ* = 0.1, significantly distort the species tree topology. The behaviors of ASTRAL were found to be consistent with theoretical deductions regarding the anomaly zones (Fig. 2e); that is, the presence of the AGT resulted in the incorrect inference of the species tree by ASTRAL. The results of our simulations suggest that a full-likelihood method, such as *BEAST, was more sensitive than ASTRAL to introgression. Even for a moderate degree of ILS at *t*_2_ = 0.9, *BEAST still estimated the false species tree (*A*, (*B, C*)) at *γ* = 0.1. There was one exception: When the time of divergence of the ghost predated the split time of the sister species for introgression from a C-derived ingroup ghost to B (Figs. S2b and c), i.e., *t*_1_ > 1.5, *BEAST delivered relatively better results. Moreover, ASTRAL and *BEAST were found to be likely to provide strong support (with local PP and PP equal to one) for the incorrect species tree topology (*A*, (*B, C*)) (Figs. 7d and 8d).

The divergence times for the MRCA of the three species A, B, and C were underestimated under all the above four introgression scenarios (Figs. 7e and 8e). For *γ* = 0 (i.e., only ILS), *BEAST accurately estimated the divergence times of the species. However, for an internal branch length of *t*_2_ ≥ 0.5, the estimates of 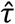 decreased sharply at *γ* = 0.1, especially with a small *t*_1_. The divergence times were estimated relative to the inferred topology. Under the same parameter settings, the correctly inferred topology ((*A, B*), *C*) showed more accurate divergence time estimates than the incorrect topology (*A*, (*B, C*)). At the same time, the results showed that the relationship between the divergence time estimates and the probability of introgression *γ* for the direction of inflow was different from that for the outflow direction. For the former, the estimated divergence times 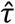 clearly decreased first and then increased to near the true value with increasing *γ* (Fig. 7e); in other words, intermediate levels of introgression tended to introduce a more pronounced bias in the estimated time. For the latter, 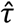 decreased sharply at first with increasing values of *γ* up to 0.4, and then stabilized close to the divergence time of the sister species A and B. (Fig. 8e). Accuracy in terms of the estimated time decreased monotonically with increasing introgression.

### Results of scenarios of Introgression between Ancestral Sister Species, and some Related Ingroup Ghost Introgressions

Figure 9 presents the results of introgression between ancestral sister species for the direction from the basal species *C* to the ancestor of the sister species. For the case of an introgression event very rapidly followed by a speciation event (say *t*_1_ = 0.05), ASTRAL performed well at *γ* < 0.5, but inferred incorrect species tree topologies (*A*, (*B, C*)) and ((*A, C*), *B*) at *γ* > 0.5 (Fig. 9b), as did *BEAST in general (Fig. 9c). The accuracy of topological inference decreased first and then increased as *γ* increased from 0.5 to 0.9 (Figs. 9b and c), which aligns very well with the results of theoretical derivation presented in Fig. 3a. For parameter combinations when the accuracy of the topological inference was low (namely, in or close to the anomaly zone), ASTRAL provided insufficient support (with a local PP of close to 0.7) and *BEAST delivered similar performance (Fig. 9d). For *t*_1_ = 0.05 and *γ* ≥ 0.4, the incorrectly inferred species tree accounted for about half of all repetitions of the simulation. *BEAST also did not provide sufficient evidence for false species tree estimates, as the PP decreased sharply from 1 to near 0.7 when the accuracy decreased. The divergence times for the MRCA of the three species A, B, and C were underestimated, and the extent of the underestimation depended on the values of *t*_2_ and *γ* (Fig. 9e). Regardless of the estimated topology of the species tree, posterior estimates for the divergence times decreased sharply with increasing *γ* for *γ* ≤ 0.4, and gradually approached the time of the introgression event.

**Figure 9.**
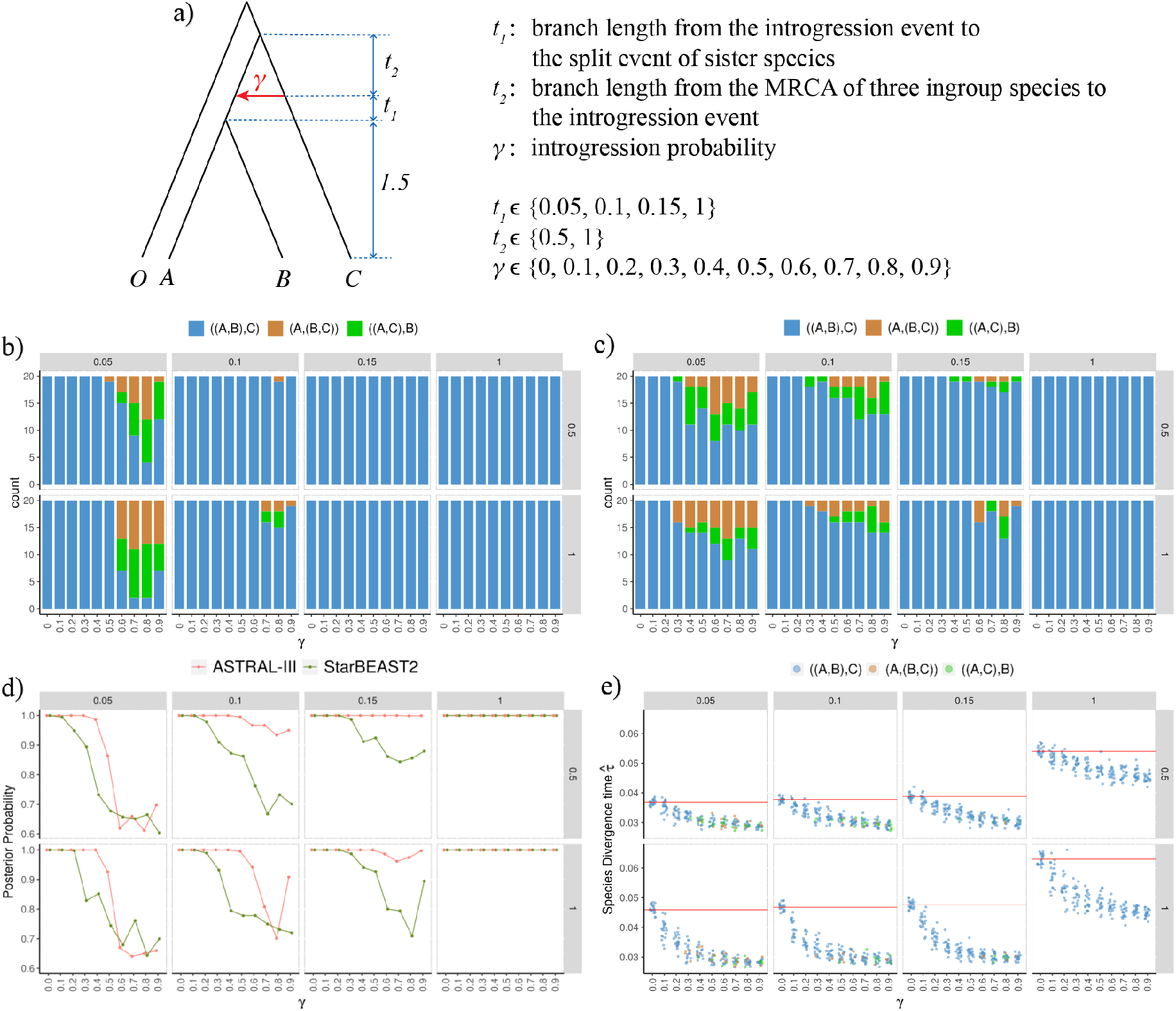
Results of simulation of introgression between an ancestral sister species and species C as donor. a) The simulated introgression scenarios and parameter settings. The outgroup species is labeled with the letter O. The lengths of the branches *t_i_* are in coalescent units. (b–e) Plots of the results of simulations. The numbers specified on the strips on the top and right of each plot represent *t*_1_ and *t*_2_, respectively, and *γ* is labeled on the x-axis. b) The numbers of topologies of three species trees inferred by ASTRAL-III (with the outgroup species O omitted for simplicity) among 20 replicates. c) The numbers of topologies of three species trees inferred by StarBEAST2. d) Local posterior probability (ASTRAL-III) and posterior clade probability (StarBEAST2) for the node of the sister species. Each data point represents the mean value across 20 replicates, and is jittered horizontally to avoid clutter. e) Estimated divergence times for the MRCA of the three ingroup species. The points and the red line represent the estimates and the true value, respectively.

In case of introgression from the ancestor of the sister species to species C (the left side of Fig. S3a), and from an ingroup ghost derived from the ancestor to C (the right side of Fig. S3a), ASTRAL inferred the species tree with 100% accuracy, and provided strong support for all parameter combinations (Figs. S3b–d), which can be attributed to the absence of AGTs in this scenario. The accuracy of *BEAST dropped slightly at large values of *γ* and *t*_1_ ≤ 0.15 (Fig. S3c), and PPs for the estimated species trees were affected by severe introgression at small values (< 0.1) of *t*_1_ (Fig. S3d). The divergence times for the MRCA of the three species were underestimated (Fig. S3e), and the tendency of these times to decline with *γ* was found to be similar to those in scenarios of introgression involving the reverse direction, i.e., *C* → ancestor.

### Results of scenarios of Introgression between Extant Sister Species, and some related Ingroup Ghost Introgressions

In scenarios of introgression between sister species and introgression from a species from the A-derived ghost to species B (Fig. S4), we fixed *t*_2_ at 0.05 for high ILS. ASTRAL inferred the species tree highly accurately, and local PPs for the estimated species trees increased with *γ*. *BEAST performed poorly for a large ILS at *γ* = 0, probably due to an insufficient number of sampled loci (50). The accuracy of topological estimation and the PP increased with the probability of introgression *γ*. The divergence times for the MRCA of the three species were little affected by introgression between extant sister species (Fig. S4e). All these results were consistent with continuous the results of a study on migration between sister species (Leache et al. 2014), which operated as a homogenizing force to reduce the discordance in the gene tree.

We also considered the scenario involving introgression from an ingroup ghost derived from the internal branch to one of the sister species (Fig. S5). Note that this scenario, in contrast to the effect of homogeneity of the two scenarios described in Figure S4, was expected to involve a reduced probability of a coalescence of sequences in the internal branch and, hence, an increased discordance in the gene tree. We set the length of the internal branch to 0.5 and moved the position of the ghost to it (*t*_2_) to change the degree of discordance of the gene tree. ASTRAL and *BEAST inferred the species tree highly accurately, regardless of the position of the ghost lineage (*t*_2_) and the probability of introgression (*γ*), and the PPs calculated by *BEAST decreased sharply as *γ* increased from 0.5 (Fig. S5d). The divergence times for the MRCA of the three species were little affected by a change in the level of introgression in this scenario (Fig. S5e).

### Results of Complex Introgression scenarios

Finally, to investigate the combined effect of two patterns of introgression on the divergence time estimates, we simulated a complex introgression scenario involving an introgression event between nonsister species and another introgression event from an outgroup ghost to species C (Fig. S6). We let *γ*_1_ = 0,0.1,0.2, and 0.3 for the introgression between nonsister species. The topology of the species tree was inferred accurately using both ASTRAL and *BEAST. For the case involving only a ghost introgression event with *γ*_1_ = 0, the divergence times for the MRCA of the three species were overestimated, as expected, with values of 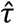 increasing with those of *γ*_2_. However, for the case of another introgression event between the nonsister species at *γ*_1_ ≥ 0.1, the divergence times were underestimated regardless of the level of introgression from the ghost (*γ*_2_), and the degree of underestimation increased with increasing *γ*_1_ (Fig. S6e).

## Discussion

Previous studies have shown that the impact of gene flow on species tree estimation can be severe in many cases (Leache et al. 2014; Solis-Lemus et al. 2016; Long and Kubatko 2018; Jiao et al. 2020). In this work, we considered a greater number and variety of gene flow scenarios under a model of episodic introgression, especially ghost introgressions that are most likely to occur in the real world. In a recent study, Tricou et al. (2021) showed that introgression between a ghost lineage and a sampled taxon can produce significant tests for introgression, but the timing, direction, and identity of lineages involved in introgression may all be inferred incorrectly when currently available approaches are used to detect introgression. Hibbins and Hahn (2021) have examined a limited number of scenarios of ghost introgression under the MSci model. In contrast to Tricou et al. (2021) and Hibbins and Hahn (2021), we are not concerned with detecting ghost introgression *per se*, but with its consequences on the inference of species tree topologies and divergence times. As an initial systematic investigation on the impact of ghost introgression on inferences of the species tree under the multispecies coalescent model, we set out to answer two general questions: First, how does ghost introgression impact estimates of the species tree topology made by using coalescent-based methods? Second, how does ghost introgression impact estimates of the divergence time obtained from full-likelihood coalescent methods? We performed this research for the simplest cases of three species, but many conclusions thus obtained appear to be applicable to more general situations by straightforward extension.

### On Estimation of Species Tree Topology

To the best of our knowledge, our work here represents the first systematic analysis of introgression derived from ghost lineages. Long and Kubatko (2018) found that introgression between ancestral sister species can lead to AGTs on three-taxon trees. We generalized this finding because anomalous behaviors were found to also occur for some scenarios of ghost introgression, including introgression from an outgroup ghost into one of the sister species or their ancestor, as well as some introgressions descending from ingroup ghost lineages. Both Solis-Lemus et al. (2016) and our study here have established the presence of AGTs on three-taxon (or unrooted AGTs on four-taxon) trees under gene flow, directly leading to inconsistencies in the estimates made by ASTRAL. By comparison, the performance of the full-likelihood method is also affected, to varying degrees, by ghost introgression (see the next two paragraphs). These results underscore the importance of paying attention to introgression from ghost lineages when conducting research in this field. When reconstructing the phylogeny for a clade, it is better to sample as many taxa as possible to minimize the risk and the resultant impact of introgression from unsampled ghost lineages.

Under many conditions of ghost introgression or introgression between extant nonsister species, coalescent-based phylogenetic methods perform poorly in estimating the history of speciation of the species because even a small amount of introgression in combination with very severe ILS can cause misleading estimates of the species tree by both ASTRAL and full-likelihood methods. Jiao et al. (2020) found that introgression between extant nonsister species can have a strong impact on species tree estimation at a high ILS, and we extended this to more cases of introgression involving ghosts, including ingroup ghost introgression (Figs. 7 and 8) and introgression from an outgroup ghost to the sister species (Fig. 6). In these scenarios, a significant interaction was noted between ILS and gene flow in the context of the possible appearance of AGTs. The higher was the level of ILS, the lower was the probability of the gene tree matching the species tree, and hence the smaller was the amount of introgression required for AGT to occur. On the other hand, even massive introgression, with more than half of the recipient genome descending from the donor lineage, may not necessarily produce AGTs. For example, in cases of introgression from an outgroup ghost to one sister species (Fig. 1b), when the length of the internal branch in the species tree is greater than the distance of the outgroup ghost (i.e., when *C*_1_ > *C*_2_), ASTRAL and full-likelihood methods can accurately infer the species tree, with a probability of introgression greater than 0.5.

The relative performance of ASTRAL and full-likelihood methods varies under different introgression scenarios. For introgression from an outgroup ghost to a sister species, the full-likelihood method was found to be more robust against gene flow than ASTRAL (Fig. 6). By contrast, in scenarios of introgression between nonsister species and some related cases of ingroup ghost introgression, ASTRAL was found to be more robust against introgression than the full-likelihood method (Figs. 7 and 8), where this is consistent with the analytical results reported by Jiao et al. (2020). This is probably because the full-likelihood method also used information retained in the branch lengths of the gene trees, in addition to data obtained from the tree topologies. The full-likelihood method is more likely to result in a unification, under the MSC model, of two species whose genes coalesced relatively early. For example, the alleles of species A and B in nonintrogressed loci coalesced first in the example shown in Figure 6a, making the full-likelihood method relatively good at estimating the true topology ((A, B), C). Conversely, the alleles of species B and C in introgressed loci coalesced first in the example shown in Figures 7a and 8a, and these scenarios rendered the full-likelihood method likely to produce a false topology (A, (B, C)).

In cases where the recipient lineage quickly split following an introgression event, coalescent-based methods were more likely to infer an incorrect species tree. This incorrect inference can be attributed to the two alleles sampled from the descendent species not having the time to coalesce, and hence perhaps frequently originating from different parental populations. This led to the prevalence of gene tree discordance and even AGTs (Solis-Lemus et al. 2016; Long and Kubatko 2018; Blair and Ané 2020). In this work, we have shown that only when the distance between the introgression event and the subsequent speciation event is shorter than 0.224 (in coalescent units) can AGTs arise. In the simple three-taxon scenarios of introgression from an outgroup ghost or basal species to the ancestor of the sister species (i.e., Figs. 5 and 9), two interesting features were identified. First, the accuracy of the topological inference was shown to decrease first and then increase with increasing probability of introgression for some combinations of branch lengths. Second, support for the inference of species tree topologies, if false, was found to be low, for both ASTRAL and the full-likelihood method. This low support can be attributed to the distribution of the topology with the same branch length for the two AGTs, and hence making it difficult to choose one of the two incorrect topologies with strong confidence.

The performance of coalescent-based methods is also affected by the direction of gene flow. For introgression involving ghost lineages, if gene flow is direction from the lineage in the species tree to the ghost, it has no way of impacting the coalescent process of alleles from the sampled species, and hence has no consequence on species tree estimation; conversely, if the ghost lineage acts as the donor of introgression, as mentioned earlier, it is possible to produce AGTs on three-taxon rooted trees, which in turn misleads species tree estimation by ASTRAL and full-likelihood methods. In cases of introgression between nonsister species, and further assuming a constant population across the species tree, inflow introgression is more likely to give rise to AGTs than outflow introgression, resulting in a lower accuracy for ASTRAL under inflow than outflow introgression (Figs. 7 and 8). In cases of introgression between ancestral sister species, if a sister lineage (*M*) split but the other (*N*) did not, introgression from *N* to *M* could produce AGTs (and, in turn, lead to biased species tree inference), but no AGTs resulted for the opposite direction of introgression from *M* to *N*.

### On estimation of Divergence Times of species

Most previous work that has used the MSC model to infer the species tree despite introgression has focused on the impact of gene flow on the inference of topology (Solis-Lemus et al. 2016; Long and Kubatko 2018; Jiao et al. 2020), but has largely ignored its effect on the estimation of divergence times. Limited exceptions include work by Leache et al. (2014) under the IM model, and Wen and Nakhleh (2018) under the MSci model. Possibly owing to the large computational cost of full-likelihood methods, the effect of gene flow on divergence time estimation has not been investigated in any systematic way, especially under the model of episodic introgression, let alone the effect of ghost introgression on the time estimates.

We found that in most scenarios of ingroup introgression, the time of root divergence among the sampled species was either underestimated or unaffected, regardless of whether ghosts were involved. This is consistent with work by Leache et al. (2014) as well as Wen and Nakhleh (2018). Introgression between nonsister species and some related ingroup ghost introgression (Figs. 7–8) generally leads to the underestimation of the time of root divergence, and involves an interaction between ILS and gene flow. For low ILS with a long internal branch in the species tree, the divergence times were underestimated severely with even modest introgression (say, *γ* = 0.1 in Figs. 7 and 8). For high ILS with a short internal branch in the species tree, the topology of the tree was more prone to be biased by the effect of introgression, but the times of root divergence were estimated accurately, and were less affected by introgression. In other words, the accurate estimation of the divergence time of species can be obtained under the prevalent ILS and the resultant, incorrect, tree topology. In case of introgression between ancestral sister species, with a long interval between the introgression event and the MRCA of the three species, the root divergence times of the three focal species were nonetheless underestimated significantly for a modest level of introgression in any direction, even though the topology of the species tree had been correctly inferred.

Most notably, introgression from an outgroup ghost generally resulted in an overestimation of the root divergence times. The introgression of the outgroup ghost brought forward the coalescence times of the introgressed alleles to the time of divergence of the ghost. Thus, it is reasonable to expect that the estimated root divergence times would increase to approach the divergence time of the ghost with increased introgression. Fortunately, compared with the underestimation caused by ingroup introgression, the magnitude of overestimation by outgroup ghost introgression was considerably weakened. Specifically, as long as the magnitude of outgroup ghost introgression was not large (*γ* < 0.5), its effect on the estimated divergence time was not significant because the coalescence times of the three alleles (one from each of the three sampled species) in nonintrogressed loci were necessarily distributed around their MRCA.

The direction of gene flow matters not only for estimating the topology of the species tree, but also for the estimation of divergence times. When introgression occurred between nonsister species, the direction of inflow or outflow made a significant difference. For the direction of inflow, the divergence time estimates were shown to decrease first and then increase to the true value with increasing intensity of introgression. For the outflow direction, the divergence time estimates were shown to decrease monotonically to the split time of the sister species (i.e., A and B) with increasing introgression. In the extreme case for both types of scenarios, introgression could be so extensive that the recipient genome would be completely replaced with the donor’s genome, necessarily resulting in the species tree (*A*, (*B, C*)) (Mallet et al. 2016). In case of such an extremity, the height of the tree root would not change for inflow introgression, but would reduce to the split time of the sister species for outflow introgression.

When introgression is a possibility during the coalescent process, it is necessary to account for both coalescence and introgression processes. A number of computationally effective methods have been successful in detecting cross-species gene flow (Patterson et al. 2012; Dalquen et al. 2017; Blischak et al. 2018; Malinsky et al. 2021). Furthermore, several phylogenetic network-based methods that extend the MSC model to accommodate introgression have been developed to simultaneously infer gene flow events and the history of speciation, including summary methods using gene tree topologies (Yu et al. 2014; Yu and Nakhleh 2015; Solis-Lemus and Ane 2016) and more effective full-likelihood methods that make direct use of sequence alignments (Wen and Nakhleh 2018; Zhang et al. 2018a; Flouri et al. 2020). We content ourselves here with noting that phylogenetic network-based methods can explicitly infer events of ghost introgression, when the length of the introgression edge is obviously large than 0. The above-mentioned programs involve expensive computation, and are not feasible for realistically sized datasets, whereas the complexity of the relevant models that simultaneously consider ILS and introgression means that large datasets with more than hundreds of loci may be necessary to obtain reliable parameter estimates. Our study implies the pressing need for the development of phylogenetic network methods that can overcome scalability issues.

## Conclusions

Cross-species introgression is common in nature, and poses challenges to species tree estimation. In the work described in this article, we systematically analyzed, for the first time, scenarios involving ghost introgression and found that it can lead to AGTs, which in turn mislead topological inference in coalescent-based methods. We also found that ingroup introgression, whether or not involving ghosts, would in general lead current species tree methods either to underestimate the divergence times for the species under investigation or not to exert any impact. However, for cases of outgroup ghost lineages acting as donors in introgression events, the divergence time was generally overestimated.

We also demonstrated that the effects of ILS and introgression on species tree estimation are closely intertwined. In many (ghost) introgression scenarios, even a weak introgression in combination with strong ILS was sufficient to lead to anomalous gene trees and, hence, biased topological inference. However, when introgression occurred between nonsister species and/or between an extant species and an ingroup ghost, the stronger the ILS was, the higher was the accuracy of estimating the root divergence times, although species tree topology was more prone to be biased.

## Acknowledgement

We would like to thank Yamei Ding for her insightful discussions.

## Funding

This work was supported by the National Key R & D Program of China (2017YFA0605104), the National Natural Science Foundation of China (31421063), the “111” Program of Introducing Talents of Discipline to Universities (B13008), and a key project of State Key Laboratory of Earth Surface Processes and Resource Ecology.

## Appendix

### Introgression from an Outgroup Ghost

For the scenario involving introgression from outgroup ghost to the ancestor of a sister species (Fig. 1b), if we assume a constant population size such that *C*_3_ = *C*_2_ + *C*_4_, we get

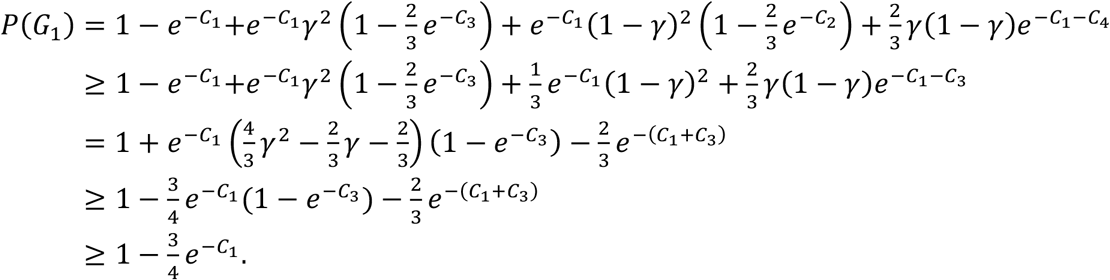

When *C*_3_ ≥ *C*_4_ and *γ* ≥ 0.5,

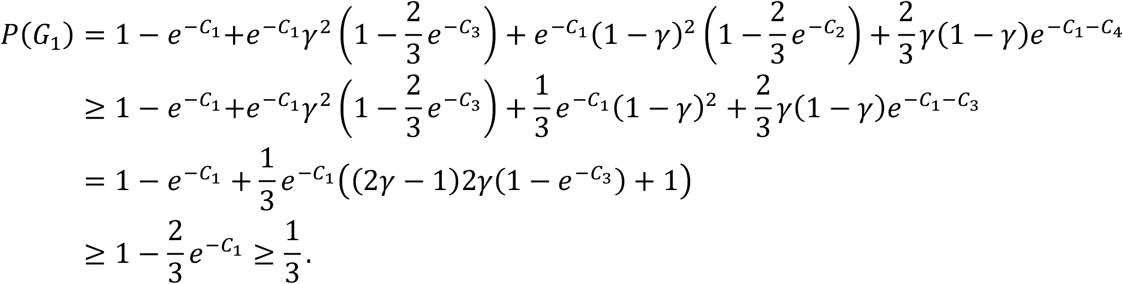

### Introgression between Ancestral Sister Species

For scenarios involving introgression from species C to the ancestor of a sister species (Fig. 3a), if we assume a constant population size such that *C*_2_ = *C*_3_,

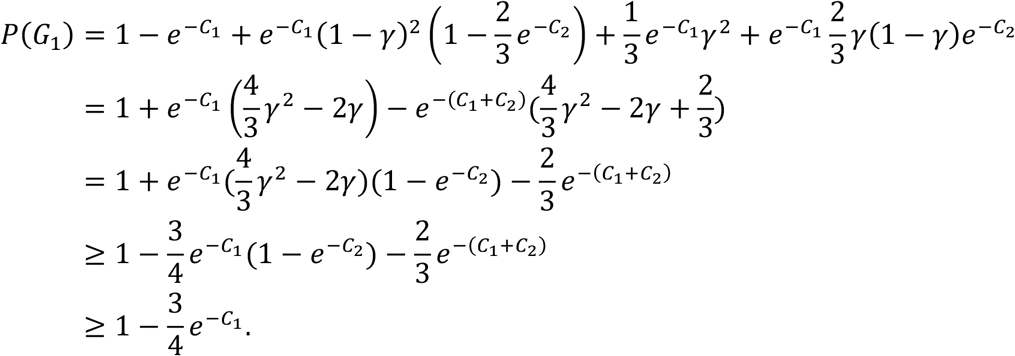

Similarly, for scenarios of introgression from a C-derived ingroup ghost to the ancestor of a sister species (Fig. 3c), if we assume a constant population size such that *C*_2_ = *C*_3_ + *C*_4_,

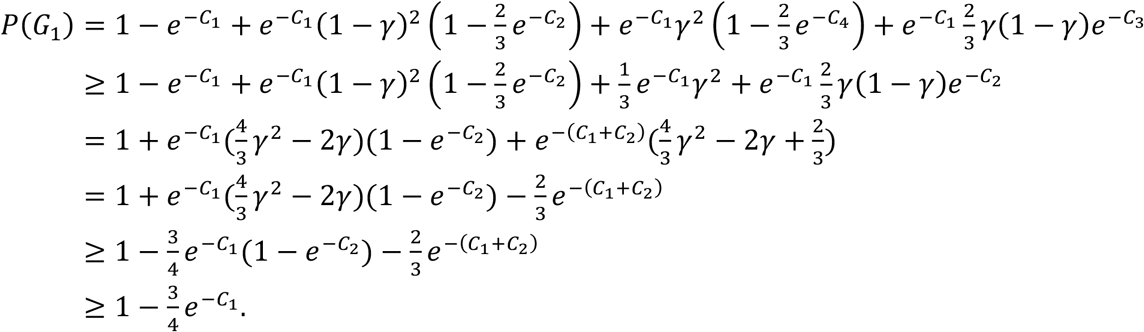

For both scenarios in Figures 3a and c, when *C*_2_ ≥ *C*_3_ and *γ* ≤ 0.5,

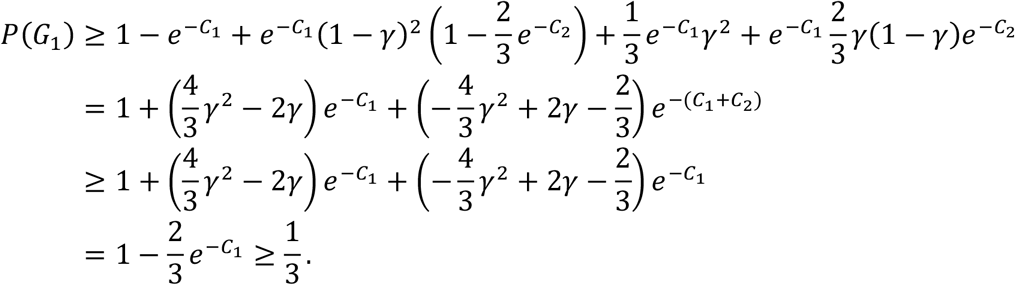

## Notes

### Competing Interest Statement

The authors have declared no competing interest.

## References

Blair C, Ané C. 2020. Phylogenetic trees and networks can serve as powerful and complementary approaches for analysis of genomic data. Syst. Biol. 69:593–601.

Blischak PD, Chifman J, Wolfe AD, Kubatko LS. 2018. HyDe: a python package for genome-scale hybridization detection. Syst. Biol. 67:821–829.

Dalquen DA, Zhu T, Yang Z. 2017. Maximum likelihood implementation of an isolation-with-migration model for three species. Syst. Biol. 66:379–398.

Degnan JH. 2013. Anomalous unrooted gene trees. Syst. Biol. 62:574–590.

Degnan JH. 2018. Modeling hybridization under the network multispecies coalescent. Syst. Biol. 67:786–799.

Degnan JH, DeGiorgio M, Bryant D, Rosenberg NA. 2009. Properties of consensus methods for inferring species trees from gene trees. Syst. Biol. 58:35–54.

Eaton DAR, Ree RH. 2013. Inferring Phylogeny and Introgression using RADseq Data: An Example from Flowering Plants (Pedicularis: Orobanchaceae). Syst. Biol. 62:689–706.

Edelman NB, Mallet J. 2021. Prevalence and Adaptive Impact of Introgression. Annu. Rev. Genet. 55:265–283.

Edwards SV, Xi Z, Janke A, Faircloth BC, McCormack JE, Glenn TC, Zhong B, Wu S, Lemmon EM, Lemmon AR, et al. 2016. Implementing and testing the multispecies coalescent model: a valuable paradigm for phylogenomics. Mol. Phylogenet. Evol. 94:447–462.

Elworth RAL, Ogilvie HA, Zhu J, Nakhleh L. 2019. Advances in computational methods for phylogenetic networks in the presence of hybridization. In: Warnow T editor. Computational Biology. New York, Springer, Cham.

Esquerré D, Keogh JS, Demangel D, Morando M, Avila LJ, Sites JW, Jr., Ferri-Yáñez F, Leaché AD. 2021. Rapid Radiation and Rampant Reticulation: Phylogenomics of South American Liolaemus Lizards. Syst. Biol. doi:10.1093/sysbio/syab058.

Flouri T, Jiao X, Rannala B, Yang Z. 2020. A Bayesian implementation of the multispecies coalescent model with introgression for phylogenomic analysis. Mol. Biol. Evol. 37:1211–1223.

Fontaine MC, Pease JB, Steele A, Waterhouse RM, Neafsey DE, Sharakhov IV, Jiang X, Hall AB, Catteruccia F, Kakani E, et al. 2015. Extensive introgression in a malaria vector species complex revealed by phylogenomics. Science 347:1258524–1258524.

Forsythe ES, Nelson ADL, Beilstein MA. 2020. Biased Gene Retention in the Face of Introgression Obscures Species Relationships. Genome Biol. Evol. 12:1646–1663.

Giarla TC, Esselstyn JA. 2015. The challenges of resolving a rapid, recent radiation: empirical and simulated phylogenomics of Philippine shrews. Syst. Biol. 64:727–740.

Heled J, Drummond AJ. 2010. Bayesian inference of species trees from multilocus data. Mol. Biol. Evol. 27:570–580.

Hey J. 2010. Isolation with migration models for more than two populations. Mol. Biol. Evol. 27:905–920.

Hibbins MS, Hahn MW. 2021. Phylogenomic approaches to detecting and characterizing introgression. Genetics doi:10.1093/genetics/iyab173.

Hudson RR. 1983. Testing the constant-rate neutral allele model with protein sequence data. Evolution 37:203–217.

Jiao X, Flouri T, Rannala B, Yang Z. 2020. The impact of cross-species gene flow on species tree estimation. Syst. Biol. 69:830–847.

Jiao X, Flouri T, Yang Z. 2021. Multispecies coalescent and its applications to infer species phylogenies and cross-species gene flow. Natl. Sci. Rev. doi:10.1093/nsr/nwab127.

Larget BR, Kotha SK, Dewey CN, Ane C. 2010. BUCKy: gene tree/species tree reconciliation with Bayesian concordance analysis. Bioinformatics 26:2910–2911.

Leache AD, Harris RB, Rannala B, Yang Z. 2014. The influence of gene flow on species tree estimation: a simulation study. Syst. Biol. 63:17–30.

Liu L, Yu L, Edwards SV. 2010. A maximum pseudo-likelihood approach for estimating species trees under the coalescent model. BMC Evol. Biol. 10:302.

Long C, Kubatko L. 2018. The effect of gene flow on coalescent-based species-tree inference. Syst. Biol. 67:770–785.

Maddison WP. 1997. Gene trees in species trees. Syst. Biol. 46:523–536.

Malinsky M, Matschiner M, Svardal H. 2021. Dsuite - fast D-statistics and related admixture evidence from VCF files. Mol Ecol Resour 21:584–595.

Mallet J, Besansky N, Hahn MW. 2016. How reticulated are species? Bioessays 38:140–149.

Mirarab S, Bayzid MS, Warnow T. 2016. Evaluating summary methods for multilocus species tree estimation in the presence of incomplete lineage sorting. Syst. Biol. 65:366–380.

Mirarab S, Nakhleh L, Warnow T. 2021. Multispecies Coalescent: Theory and Applications in Phylogenetics. Annu Rev Ecol Evol Syst 52:247–268.

Mirarab S, Reaz R, Bayzid MS, Zimmermann T, Swenson MS, Warnow T. 2014. ASTRAL: genome-scale coalescent-based species tree estimation. Bioinformatics 30:541–548.

Molloy EK, Warnow T. 2018. To include or not to include: the impact of gene filtering on species tree estimation methods. Syst. Biol. 67:285–303.

Nichols R. 2001. Gene trees and species trees are not the same. Trends Ecol. Evol. 16:358–364.

Ogilvie HA, Bouckaert RR, Drummond AJ. 2017. StarBEAST2 brings faster species tree inference and accurate estimates of substitution rates. Mol. Biol. Evol. 34:2101–2114.

Pamilo P, Nei M. 1988. Relationships between gene trees and species trees. Mol. Biol. Evol. 5:568–583.

Patterson N, Moorjani P, Luo Y, Mallick S, Rohland N, Zhan Y, Genschoreck T, Webster T, Reich D. 2012. Ancient Admixture in Human History. Genetics 192:1065–1093.

Pease JB, Hahn MW. 2015. Detection and polarization of introgression in a five-taxon phylogeny. Syst. Biol. 64:651–662.

Pierron D, Heiske M, Razafindrazaka H, Rakoto I, Rabetokotany N, Ravololomanga B, Rakotozafy LM-A, Rakotomalala MM, Razafiarivony M, Rasoarifetra B, et al. 2017. Genomic landscape of human diversity across Madagascar. Proc. Natl. Acad. Sci. USA 114:E6498–E6506.

Rambaut A, Grass NC. 1997. Seq-Gen: an application for the Monte Carlo simulation of DNA sequence evolution along phylogenetic trees. Bioinformatics 13:235–238.

Rannala B, Yang Z. 2003. Bayes estimation of species divergence times and ancestral population sizes using DNA sequences from multiple loci. Genetics 164(4):1645–1656.

Rannala B, Yang Z. 2017. Efficient Bayesian species tree inference under the multispecies coalescent. Syst. Biol. 66:823–842.

Sayyari E, Mirarab S. 2016. Fast coalescent-based computation of local branch support from quartet frequencies. Mol. Biol. Evol. 33:1654–1668.

Slatkin M, Maddison WP. 1989. A cladistic measure of gene flow inferred from the phylogenies of alleles. Genetics 123:603–613.

Smith O, Nicholson WV, Kistler L, Mace E, Clapham A, Rose P, Stevens C, Ware R, Samavedam S, Barker G, et al. 2019. A domestication history of dynamic adaptation and genomic deterioration in Sorghum. Nat. Plants 5:369–379.

Solis-Lemus C, Ane C. 2016. Inferring phylogenetic networks with maximum pseudolikelihood under incomplete lineage sorting. PLoS Genet. 12:e1005896.

Solis-Lemus C, Yang M, Ane C. 2016. Inconsistency of species tree methods under gene flow. Syst. Biol. 65:843–851.

Taylor SA, Larson EL. 2019. Insights from genomes into the evolutionary importance and prevalence of hybridization in nature. Nat. Ecol. Evol. 3:170–177.

Telford MJ, Budd GE, Philippe H. 2015. Phylogenomic insights into animal evolution. Curr. Biol. 25:876–887.

Thawornwattana Y, Dalquen D, Yang Z. 2018. Coalescent Analysis of Phylogenomic Data Confidently Resolves the Species Relationships in the *Anopheles gambiae* Species Complex. Mol. Biol. Evol. 35:2512–2527.

Tiley GP, Poelstra JW, Dos Reis M, Yang Z, Yoder AD. 2020. Molecular clocks without rocks: new solutions for old problems. Trends Genet. 36:845–856.

Tricou T, Tannier E, de Vienne DM. 2021. Ghost lineages highly influence the interpretation of introgression tests. bioRxiv doi:10.1101/2021.03.30.437672.

Wen D, Nakhleh L. 2018. Coestimating reticulate phylogenies and gene trees from multilocus sequence data. Syst. Biol. 67:439–457.

Wu D-D, Ding X-D, Wang S, Wójcik JM, Zhang Y, Tokarska M, Li Y, Wang M-S, Faruque O, Nielsen R, et al. 2018. Pervasive introgression facilitated domestication and adaptation in the Bos species complex. Nat. Ecol. Evol. 2:1139–1145.

Yin J, Zhang C, Mirarab S. 2019. ASTRAL-MP: scaling ASTRAL to very large datasets using randomization and parallelization. Bioinformatics 35:3961–3969.

Yu Y, Degnan JH, Nakhleh L. 2012. The probability of a gene tree topology within a phylogenetic network with applications to hybridization detection. PLoS Genet. 8:456–465.

Yu Y, Dong J, Liu KJ, Nakhleh L. 2014. Maximum likelihood inference of reticulate evolutionary histories. Proc. Natl. Acad. Sci. USA 111:16448–16453.

Yu Y, Nakhleh L. 2015. A maximum pseudo-likelihood approach for phylogenetic networks. BMC Genom. 16:S10.

Zhang C, Ogilvie HA, Drummond AJ, Stadler T. 2018a. Bayesian inference of species networks from multilocus sequence data. Mol. Biol. Evol. 35:504–517.

Zhang C, Rabiee M, Sayyari E, Mirarab S. 2018b. ASTRAL-III: polynomial time species tree reconstruction from partially resolved gene trees. BMC Bioinform. 19:153.

Zhang D, Rheindt FE, She H, Cheng Y, Song G, Jia C, Qu Y, Alström P, Lei F. 2021. Most Genomic Loci Misrepresent the Phylogeny of an Avian Radiation Because of Ancient Gene Flow. Syst. Biol. 70:961–975.

