## Supplementary figures1-6 for "Impact of Ghost Introgression on Coalescent-based Species Tree Inference and Estimation of Divergence Time"

### SUPPLEMENTARY MATERIAL

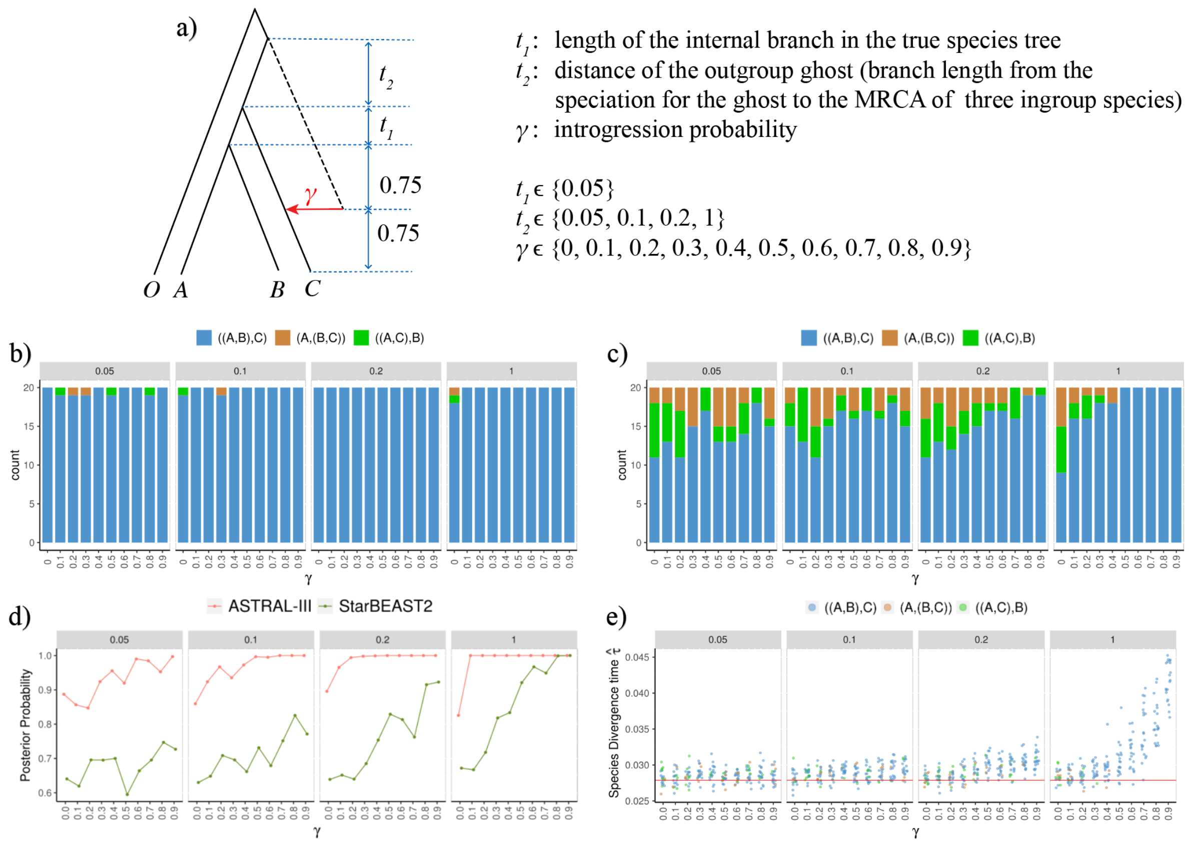


FIGURE S1. Results of simulation of introgression from an outgroup ghost to species C. a) The simulated introgression scenarios and parameter settings. The outgroup species is labeled with the letter O. The lengths of the branches $t_i$ are in coalescent units. (b–e) Plots of the results of simulations. The numbers specified on the strips on the top of each plot represent $t_2$, and $\gamma$ is labeled on the x-axis. b) The numbers of topologies of three species trees inferred by ASTRAL-III (with the outgroup species O omitted for simplicity). c) The numbers of topologies of three species trees inferred by StarBEAST2. d) Local posterior probability (ASTRAL-III) and posterior clade probability (StarBEAST2) for the node of the sister species. Each data point represents the mean value across 20 replicates, and is jittered horizontally to avoid clutter. e) Estimated divergence times for the MRCA of the three ingroup species. The points and the red line represent the estimates and the true value, respectively.

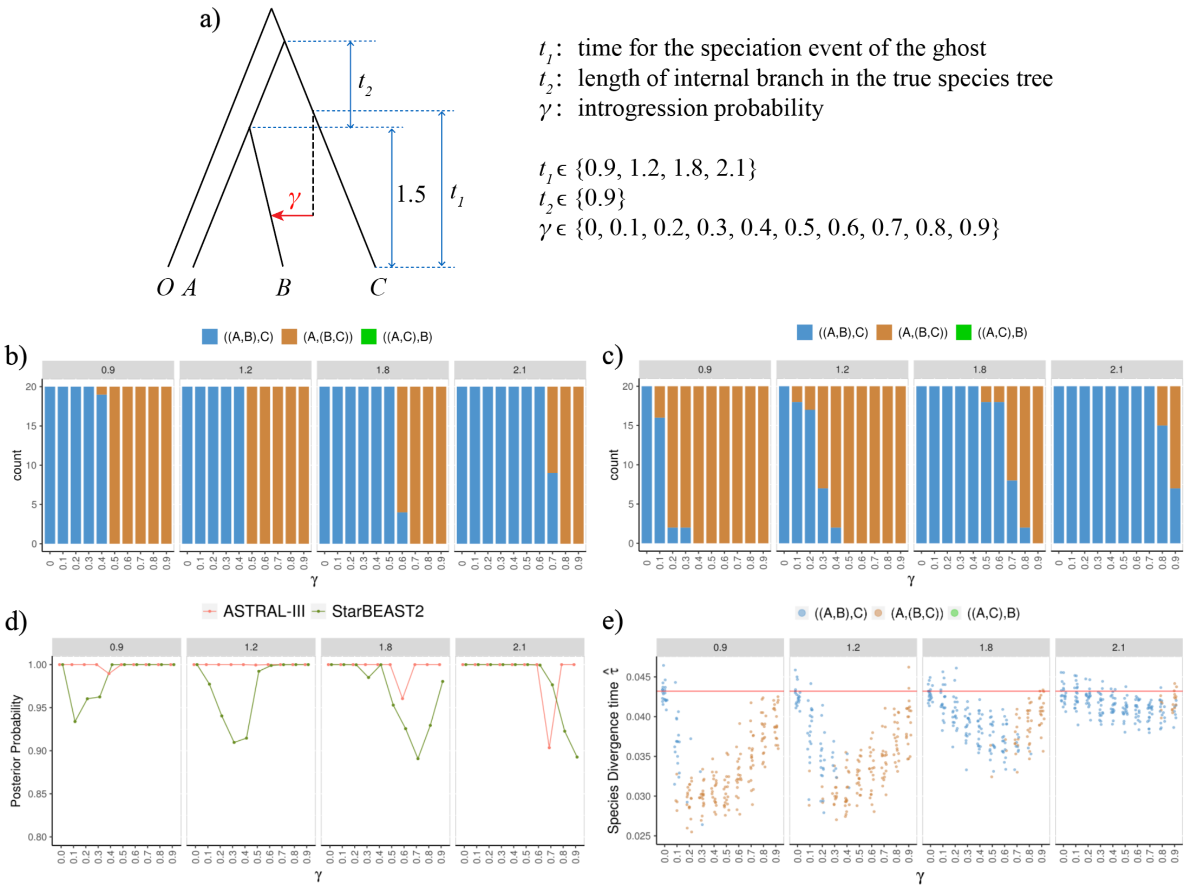


FIGURE S2. Results of simulation of introgression from a C-derived ingroup ghost to B. a) The simulated introgression scenarios and parameter settings. The outgroup species in each case is labeled with O. The lengths of the branches $t_i$ are in coalescent units. (b–e) Plots of the results of simulations. The numbers specified on the strips on the top represent $t_1$, and $\gamma$ is labeled on the x-axis. b) The numbers of topologies of three species trees inferred by ASTRAL-III (with the outgroup species O omitted for simplicity) among 20 replicates. c) The numbers of topologies of three species trees inferred by StarBEAST2. d) Local posterior probability (ASTRAL-III) and posterior clade probability (StarBEAST2) for the node of the sister species. Each data point represents the mean value across 20 replicates, and is jittered horizontally to avoid clutter. e) Estimated divergence times for the MRCA of the three species. The points and the red line represent the estimates and the true value, respectively.

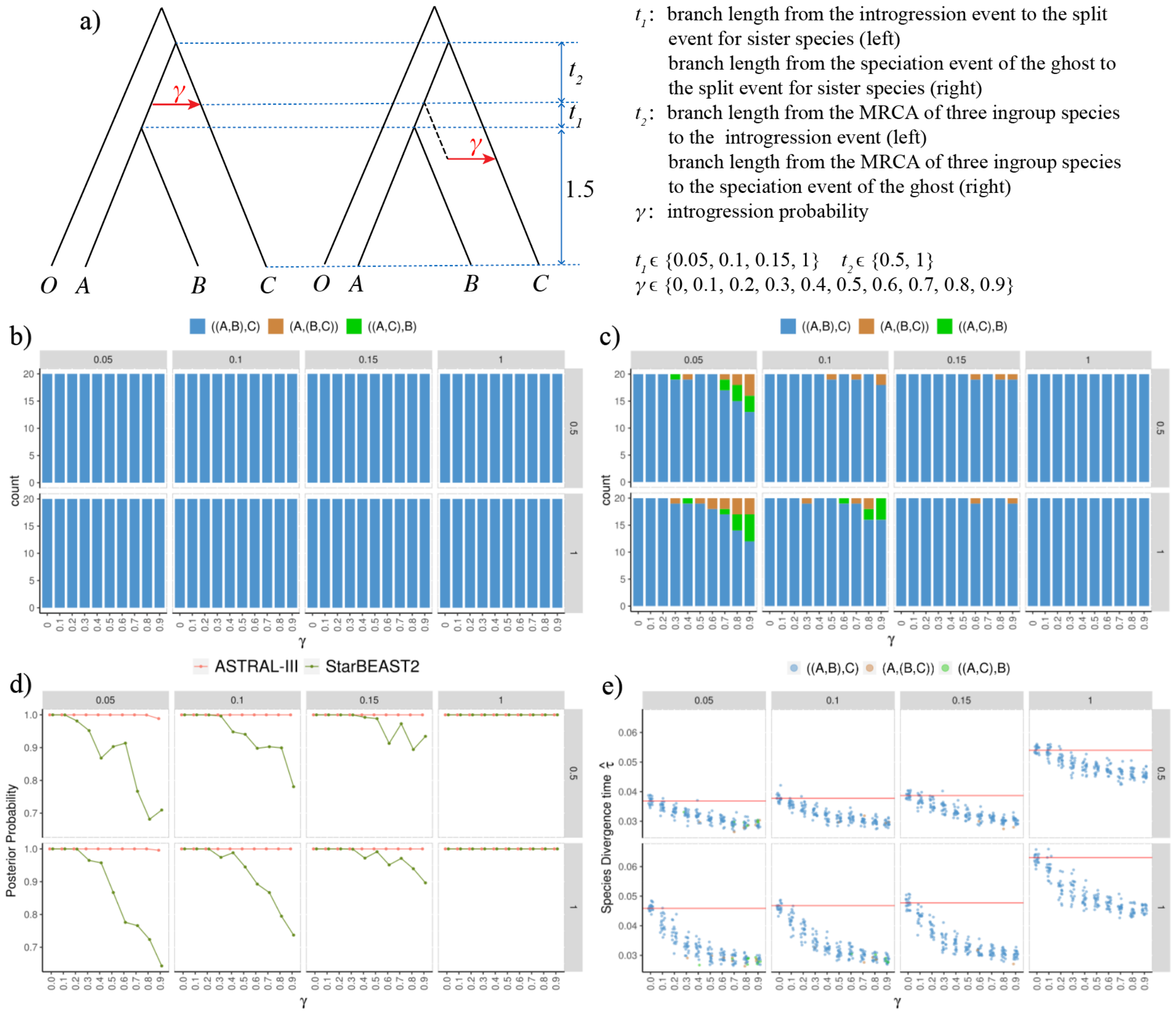


FIGURE S3. Results of simulation of introgression between ancestral sister species with species C as the recipient, or introgression from an ingroup ghost derived from the ancestor to species C. a) The simulated introgression scenarios and parameter settings. The outgroup species in each case is labeled with the letter O. The lengths of the branches $t_i$ are in coalescent units. The two scenarios generated the same distribution of gene trees with coalescent times under the same parameter settings $(t_1, t_2, \gamma)$ when one individual was sampled per species, and thus had identical results in our simulation. (b–e) Plots of the results of simulations. The numbers specified on the strips on the top and right of each plot represent $t_1$ and $t_2$, respectively, and $\gamma$ is labeled on the x-axis. b) The numbers of topologies of three species trees inferred by ASTRAL-III (with the outgroup species O omitted for simplicity). c) The numbers of topologies of three species trees inferred by StarBEAST2. d) Local posterior probability (ASTRAL-III) and posterior clade probability (StarBEAST2) for the node of the sister species. Each data point represents the mean value across 20 replicates, and is jittered horizontally to avoid clutter. e) Estimated divergence times for the MRCA of the three ingroup species. The points and the red line represent the estimates and the true value, respectively.

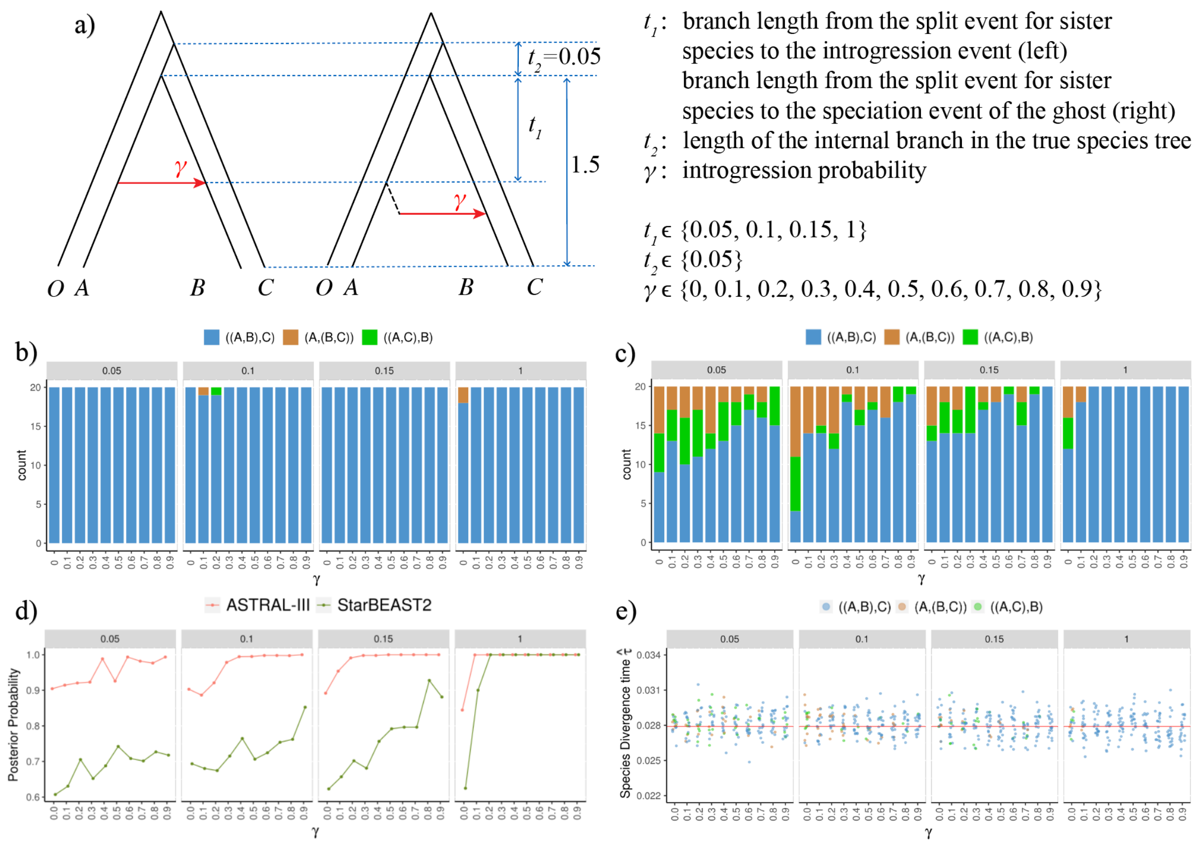


FIGURE S4. Results of simulation of introgression between sister species, or introgression from an A-derived ingroup ghost to B. a) The simulated introgression scenarios and parameter settings. The outgroup species in each case is labeled with the letter O. The lengths of the branches $t_i$ are in coalescent units. The two scenarios generated the same distribution of gene trees with coalescent times under the same parameter settings ($t_1$,$t_2$,$\gamma$) when one individual was sampled per species, and thus had identical results in our simulation. (b–e) Plots of the results of simulations. The numbers specified on the strips on the top of each plot represent $t_1$, and $\gamma$ is labeled on the x-axis. b) The numbers of topologies of three species trees inferred by ASTRAL-III (with the outgroup species O omitted for simplicity). c) The numbers of topologies of three species trees inferred by StarBEAST2. d) Local posterior probability (ASTRAL-III) and posterior clade probability (StarBEAST2) for the node of the sister species. Each data point represents the mean value across 20 replicates, and is jittered horizontally to avoid clutter. e) Estimated divergence times for the most recent common ancestor of the three ingroup species. The points and the red line represent the estimates and the true value, respectively.

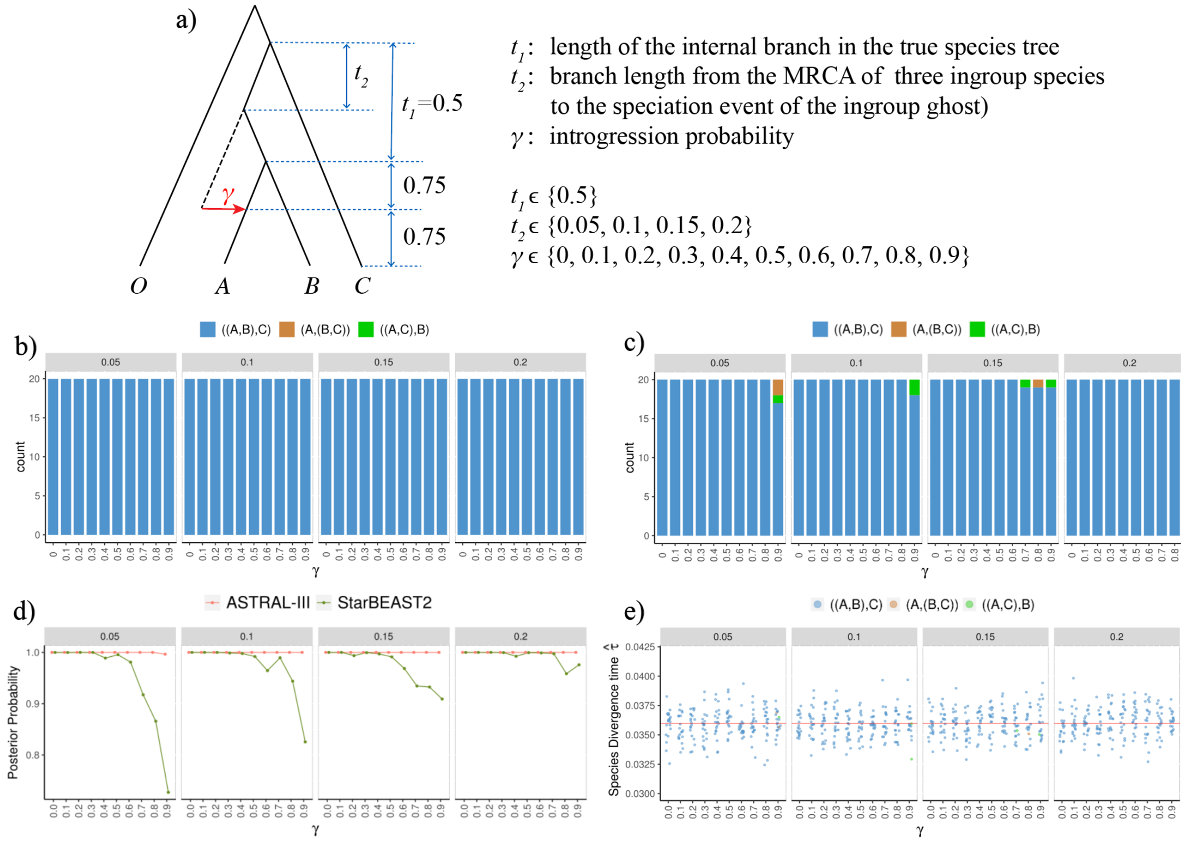


FIGURE S5. Results of simulation of introgression from an ingroup ghost, derived from the ancestor of sister species, to one of the sister species A. a) The simulated introgression scenarios and parameter settings. The outgroup species is labeled with the letter O. The lengths of the branches $t_i$ are in coalescent units. (b–e) Plots of the results of simulations. The numbers specified on the strips on the top of each plot represent $t_2$ and $\gamma$ is labeled on the x-axis. b) The numbers of topologies of three species trees inferred by ASTRAL-III (with the outgroup species O omitted for simplicity). c) The numbers of topologies of three species trees inferred by StarBEAST2. d) Local posterior probability (ASTRAL-III) and posterior clade probability (StarBEAST2) for the node of the sister species. The point represents the mean value across 20 replicates, and is jittered horizontally to avoid clutter. e) Estimated divergence times for the MRCA of the three ingroup species. The points and the red line represent the estimates and the true value, respectively.

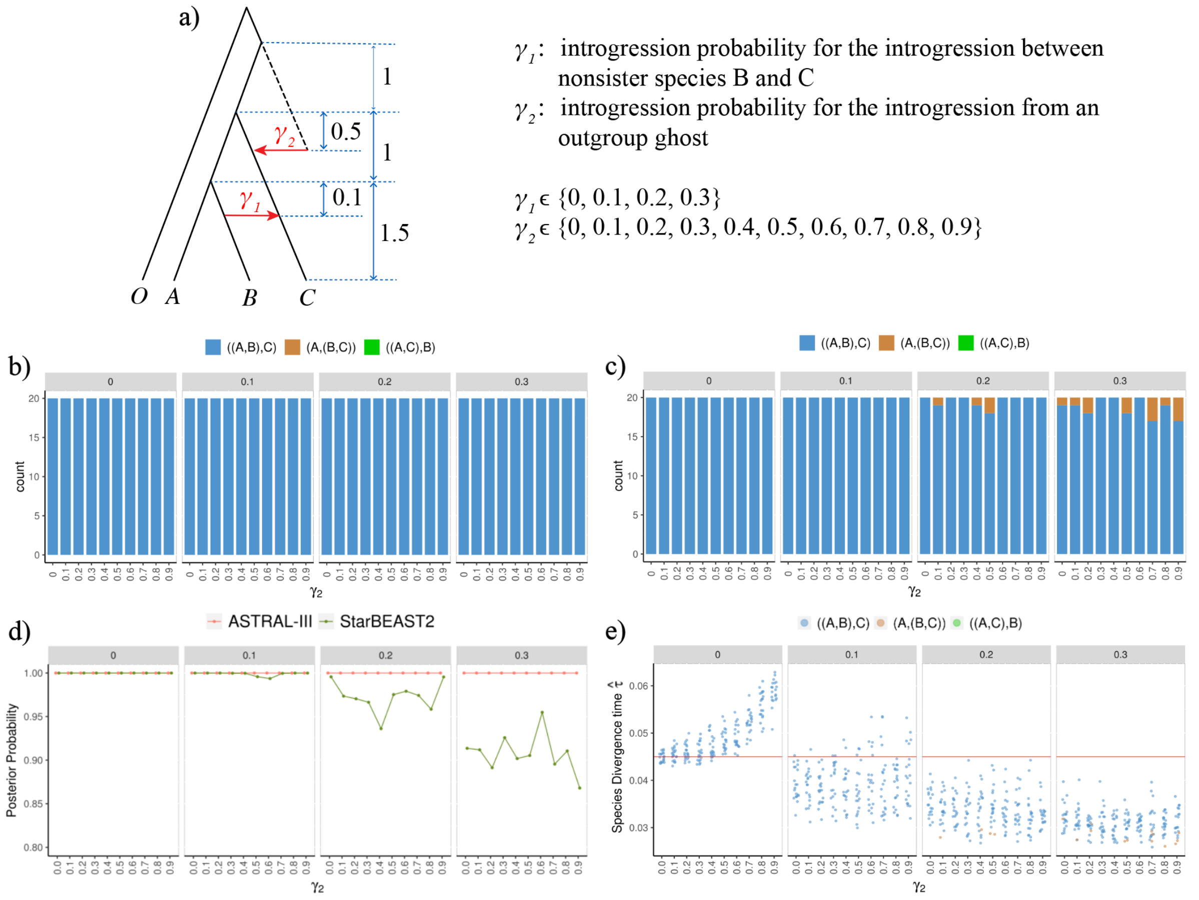


FIGURE S6. Results of simulation of complex introgression scenarios. a) The simulated introgression scenarios and parameter settings. The outgroup species is labeled with the letter O. The lengths of the branches are in coalescent units. (b–e) Plots of the results of simulations. The numbers specified on the strips on the top of each plot represent $\gamma_1$, and $\gamma_2$ is labeled on the x-axis. b) The numbers of topologies of three species trees inferred by ASTRAL-III (with the outgroup species O omitted for simplicity). c) The numbers of topologies of three species trees inferred by StarBEAST2. d) Local posterior probability (ASTRAL-III) and posterior clade probability (StarBEAST2) for the node of the sister species. Each data point represents the mean value across 20 replicates, and is jittered horizontally to avoid clutter. e) Estimated divergence times for the most recent common ancestor of the three ingroup species. The points and the red line represent the estimates and the true value, respectively.
